# PDZD-8 interacts with LGG-2/LC3 to promote autolysosome formation

**DOI:** 10.64898/2026.01.26.701500

**Authors:** Alexandre Pouget, Valentin Meire, Céline Largeau, Claire Boulogne, Baptiste Roelens, Diego Javier Zea, Christophe Lefebvre, Renaud Legouis, Emmanuel Culetto

## Abstract

LGG-2/LC3 and LGG-1/GABARAP are members of the Atg8 ubiquitin-like protein family, playing central roles in autophagy. Through a yeast two-hybrid screen, we identified the endoplasmic reticulum protein PDZD-8 as a novel interactor of the *C. elegans* autophagy protein LGG-2/LC3. We demonstrate that PDZD-8 binds specifically to LGG-2 and not LGG-1 via a canonical LC3-interacting region (LIR) and colocalizes with LGG-2-positive autophagosomes *in vivo*. Loss of PDZD-8 leads to a transient accumulation of enlarged LGG-2-positive autophagosomes that remain closely associated with lysosomes. Genetic disruption of core autophagy genes suppresses the *pdzd-8* mutant phenotype, indicating that this phenotype is dependent on autophagy. Moreover, loss of *rab-7*, which blocks autophagosome-lysosome fusion, also suppresses the *pdzd-8* phenotype, suggesting that the RAB-7-dependent fusion step is epistatic to PDZD-8 and that PDZD-8 acts downstream of autophagosome–lysosome maturation. Microscopy approaches (TEM, CLEM, and FIB-SEM) reveal an accumulation of fused autophagosome-lysosome structures, pointing to a transient block in autophagic flux in *pdzd-8* mutants. Furthermore, PDZD-8 lacking either the SMP lipid-transfer domain or the LIR motif phenocopies the *pdzd-8* null mutant, underscoring the importance of both domains. Our findings support a model in which PDZD-8, in conjunction with LGG-2/LC3, mediates ER-autophagosome/autolysosome contacts that are critical for subsequent autolysosome completion.

**Significance Statement:** Autophagy is a conserved cellular degradation pathway essential for maintaining homeostasis and responding to stress. Although the molecular machinery driving autophagosome formation and fusion with lysosomes is relatively well-characterized, the contribution of membrane contact sites to autophagic flux remains poorly understood. Here, we identify PDZD-8, an endoplasmic reticulum (ER) protein, as a novel interactor of the *Caenorhabditis elegans* LC3/Atg8 homolog LGG-2. We show that PDZD-8 regulates the completion of autolysosome maturation by bridging ER-autophagosome/autolysosome contacts. Loss of PDZD-8 results in the accumulation of enlarged autophagosomes adjacent to lysosomes and a transient block in autophagic flux, a phenotype that depends on the RAB-7-mediated tethering step. These findings reveal a previously unrecognized role for PDZD-8 in coordinating ER-autophagosome/autolysosome dynamics during autophagy.

## Introduction

The lysosome is the central degradative organelle of eukaryotic cells, responsible for breaking down and recycling a broad range of macromolecules and cytoplasmic cargoes. It functions as a central hub that coordinates anabolic and catabolic programs through nutrient availability and stress signaling. Under nutrient-rich conditions, motile lysosomes enriched in PI3P support mTORC1-dependent anabolic signaling, whereas nutrient deprivation induces a transition toward immobile, PI4P-enriched lysosomes in which mTORC1 is inactive and degradative capacity is enhanced (1, 2). The delivery of cargo to lysosomes occurs through several pathways, including autophagy and endocytosis, thereby ensuring cellular homeostasis and cell viability across varying environmental conditions (3). Within the lysosomal lumen, acidic hydrolases degrade proteins, lipids, and organelles into reusable metabolites. The efficient lysosomal degradation critically depends on membrane dynamics, inter-organelle communication, and fusion with cargo-delivering compartments (4).

Autophagy encompasses several conserved lysosome-dependent degradation pathways that are essential for cellular adaptation to stress and the maintenance of homeostasis. Cargoes can be directly addressed to the lysosomes by microautophagy and chaperone-mediated autophagy, or through the formation of a double-membrane organelle, termed an autophagosome, that engulfs cytoplasmic material. Autophagosomes subsequently fuse with lysosomes to generate autolysosomes, where cargo degradation occurs (5). This macroautophagy (hereafter referred to as autophagy) process is tightly regulated to allow fusion when autophagosomes are closed and requires the coordinated action of Rab GTPases, tethering factors, and SNARE complexes (6). The presence of the ubiquitin-like Atg8 proteins at the autophagosomal membrane is also crucial for fusion; however, the specificity of the seven LC3/GABARAP members is complex and not well understood in mammals (7). Phosphoinositide remodeling and PI4P production have emerged as key regulators in the late steps of autophagy. In mammalian cells, GABARAP recruits phosphatidylinositol 4-kinase 2A (PI4K2A) to autophagosomal membranes, promoting local PI4P production, which is required for efficient autophagosome–lysosome fusion (8). Both PI4K2A and PI4P itself scaffold the SNARE proteins STX17 and SNAP29 to facilitate docking and fusion. Studies on the nematode *Caenorhabditis elegans*, possessing only one LC3 (LGG-2) and one GABARAP (LGG-1) homologue, identified a function for LGG-2 for autolysosome maturation during developmental mitophagy (9).

Perturbations in membrane lipid composition impair autophagosome–lysosome contact formation and fusion dynamics, thereby limiting degradative capacity (10). Recently, the lipid transfer protein PDZD8 emerged as a regulator of lysosomal homeostasis at the interface between membrane organization and degradative pathways. PDZD8 is an ER membrane protein, a paralog of the yeast ERMES component Mmm1, which mediates phospholipid trafficking at ER-mitochondria contact sites (11). In metazoans, PDZD8 functions as a tether at ER-late endosome/lysosome contact sites through its interaction with Rab7, enabling lipid transfer between these organelles (12–15). PDZD8 also participates in ER-mitochondria tethering to regulate endolysosomal positioning, and is essential for neuronal polarity and outgrowth (16–18). Consistent with these roles, PDZD8 dysfunction leads to cognitive defects in humans, mice, and *Drosophila*. In *C. elegans*, PDZD-8 cooperates with the SMP protein TEX-2 to regulate phosphoinositide PI(4,5)P2 transport during embryogenesis between the endosome and ER (19). During osmotic stress, PDZD8 was shown to mediate lysosomal membrane expansion by allowing lipid transfer from the ER at ER-lysosome contact sites, thereby promoting lysosomal integrity and cell survival (20).

Recent studies have suggested a link between autophagy and PDZD8 functions (21, 22), but no direct interaction with the autophagy machinery has been documented. Here, we identify the *C. elegans* PDZD-8 as a novel interactor of the autophagy protein LGG-2/LC3 and demonstrate that this interaction contributes to autolysosome formation in heat stress conditions. Our findings reveal a novel molecular connection between the ER lipid transfer machinery and the autophagosomal membrane, supporting a specific contact site involved in autophagosome maturation.

## Results

### PDZD-8 interacts and partially colocalizes with LGG-2

We previously performed yeast two-hybrid (Y2H) screens to identify the differential interactomes of the two homologues of the ubiquitin-like protein Atg8, LGG-2/LC3 and LGG-1/GABARAP (9, 23). The LGG-2 bait identified 37 cDNA clones encoding the PDZ domain containing 8 (PDZD-8) protein (Fig. 1*A*). PDZD-8 is a multi-domain protein, which contains SMP, PDZ, C1, and CC domains, anchored to the ER membrane via its transmembrane domain (Fig. 1*A*, and SI Appendix, Fig S1*A*)(12, 13, 15, 16, 18). The PDZ and C1 domains are separated by a large non-conserved intrinsically disordered region, which in mammal cells enables PDZD8 to form condensates at the surface of the endoplasmic reticulum (24). The *pdzd-8* locus is a genetically complex locus that encodes 11 distinct isoforms through alternative transcript variants (SI Appendix, Fig S1*B* and supporting text 1).

**Figure 1:**
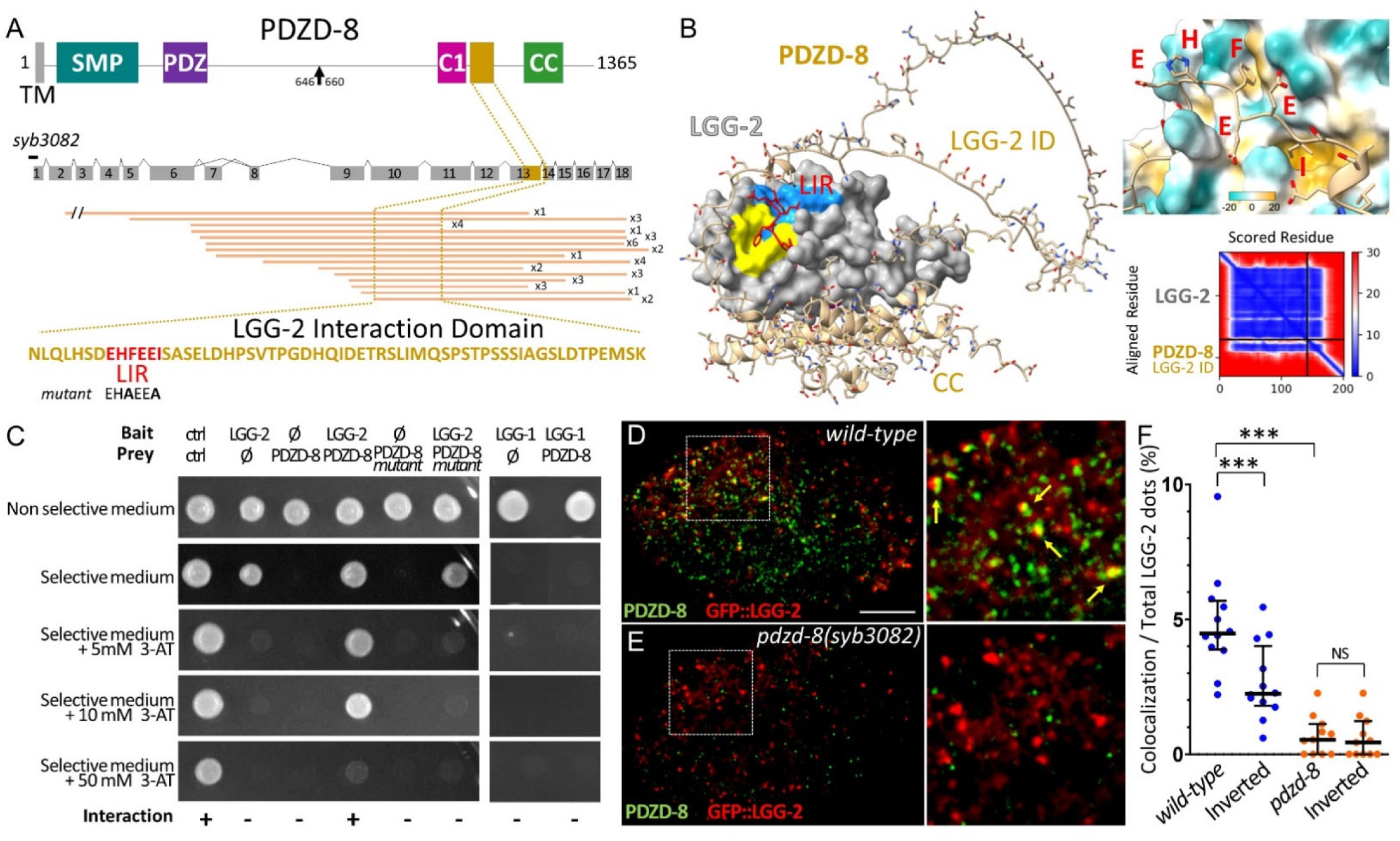
PDZD-8 directly interacts and partially colocalizes with LGG-2. (*A*) Schematic representation of the PDZD-8 protein and locus.TM (Transmembrane), SMP (Synaptotagmin-like mitochondrial-lipid-binding), PDZ (PSD95/Dlg/ZO1) C1, CC (coiled coil). The brown rectangle delineates the LGG-2 interaction domain containing an LC3-interacting Region (LIR). The black arrow indicates the epitope for PDZD-8 antibodies. Numbered grey boxes represent the exons and *syb3082* the null allele containing a small deletion and frameshift. Horizontal orange lines are cDNA clones identified in the yeast two-hybrid screen. (*B*) AlphaFold2 prediction model of interaction between PDZD-8 and LGG-2, with its predicted aligned error plot. Hydrophobic W-site (yellow) and L-site (blue) pockets are magnified in the inset, with light blue and brown representing polar and hydrophobic amino acids, respectively. (*C*) Yeast two-hybrid assays showing that LGG-2 but not LGG-1 interacts with the PDZD-8[LGG-2-ID] through its LIR motif. The inhibitor 3-aminotriazole (3AT) is used to assess the strength of the interactions and block the auto-activation of LGG-2. (*D-F*) Single merged confocal images of a GFP::LGG-2 embryo, immunostained with antiPDZD-8 (green) and antiGFP (red) antibodies in wild type (*D*) and *pdzd-8(syb3082)* mutant (*E*). The right panels are a magnified view of the box regions with colocalization events indicated by yellow arrows. Scale bar is 10µm. (*F*) Quantification of colocalization based on the distance between centers of mass. Inverted indicates the random colocalization after the original red channel was rotated to 180°. n=12,11, Bars indicate the median, the 25th and 75th percentiles. Kruskal-Wallis test with Bonferroni correction *** p<0.001, NS non-significant. See also SI-Figure S1, SI-Figure S2.

The minimal overlap between the cDNA clones defined a 59-amino-acid region that was named the LGG-2 interacting domain [LGG-2-ID] of PDZD-8 (Fig. 1*A*). The iLIR analysis (https://ilir.warwick.ac.uk/) (42) of the PDZD-8[LGG-2-ID] revealed one high-score archetypal LC3-interacting region (LIR) DEH<u>F</u>EE<u>I</u>. The protein structure prediction tool AlphaFold2 (AF2) (25) was then used to simulate the interaction between LGG-2, whose crystal structure is known (26). The average pLDDT score of AF2-predicted model for PDZD-8[LGG-2-ID] is low (52.42), suggesting an intrinsically disordered protein region. Still, the LIR was predicted to interact with a high confidence score (ipTM score 0.74) with the W and L hydrophobic pockets of LGG-2 (Fig. 1*B*, and *SI Appendix*, Fig.S1*C*).

To check the specificity of the interaction, we performed a one-by-one Y2H assay using either LGG-1 or LGG-2 as baits and the PDZD-8[LGG-2-ID] as prey (Fig. 1*C* and *SI Appendix*, Fig.S1*D*). The assay confirmed that PDZD-8[LGG-2-ID] specifically interacts with LGG-2 but not LGG-1, moreover, introducing two point-mutations in the LIR (EH**A**EE**A**) completely abolished this interaction (Fig. 1*C*). These data indicated that PDZD-8 could specifically interact with LGG-2 through its LIR domain.

To confirm this interaction *in vivo,* we generated a polyclonal antibody against the PDZD-8 protein, and we produced the *pdzd-8(syb3082)* null mutant by CRISPR-Cas9 (Fig. 1*A* and *SI Appendix*, Fig. S2). Immunofluorescent analyses with this antibody confirmed that PDZD-8 localizes as puncta at the ER in the embryo (SI Appendix, Fig. S2 *E-F’’*), similarly to the PDZD-8::mNeonGreen pattern (mNG, supplementary Fig. S2 *G-J*) (19). An immunofluorescent analysis of PDZD-8 and GFP::LGG-2 in the embryo revealed a low but significant colocalization, which was confirmed by visualizing endogenous LGG-2 and the PDZD-8::mNeonGreen (PDZD-8::mNG) (Fig. 1*D* and *F*, and *SI Appendix*, Fig. S1*F*). The co-localization events are rare but not fortuitous as revealed by the quantification of the *pdzd-8* null mutant (Fig. 1*E*) or after the inversion of one channel image (Fig. 1*F*, and *SI Appendix*, Fig. S1*E* *and* *G*). This result indicated that a subset of PDZD-8 colocalizes with LGG-2-positive vesicles *in vivo*, supporting a likely direct interaction between the two proteins. Together these data suggested that PDZD-8 through its interaction with LGG-2 could be involved in autophagy.

### *pdzd-8* inactivation affects stress-induced autophagy

The *pdzd-8 (syb3082)* mutant exhibited no obvious developmental or morphological phenotype and a lifespan similar to that of wild-type animals (*SI Appendix*, Fig. S3*A-E*), indicating that PDZD-8 is not essential under non-stress conditions. To investigate PDZD-8 functions in autophagy, we first characterized its expression pattern (Fig. 2*A* and *SI Appendix*, Fig. S2) during development using IF or PDZD-8::mNG. Despite a low level of expression, the PDZD-8 pattern was characteristic of an ER protein, with localization around the nucleus, around the centrosome, being part of the centriculum (27), as well as in the intermediate and peripheral ER. However, its punctate pattern indicated that PDZD-8 localized to subdomains of the ER. PDZD-8 is ubiquitously expressed in the early embryo, but it is weaker in the P1 compared to the AB lineage. In larvae and adults, PDZD-8 was also very weakly expressed but detectable in the epidermis and the nervous system (Fig. 2*A* and *SI Appendix*, Fig. S2*K-L’*) which is consistent with its embryonic expression in AB lineage.

**Figure 2:**
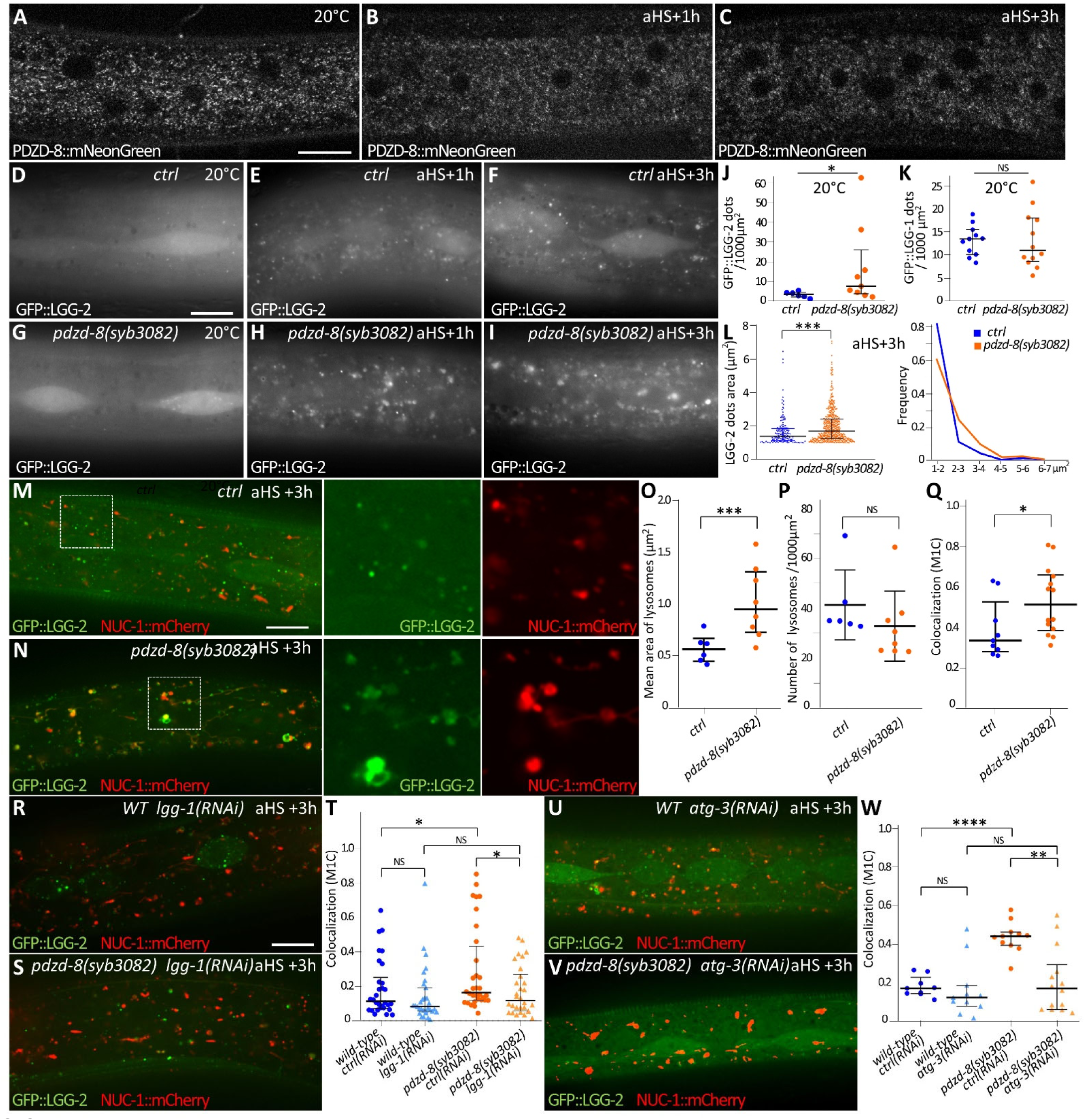
*pdzd-8* mutant is defective in stress-induced autophagy. (*A-C*) Confocal images of PDZD-8::mNeonGreen in the epidermis of L4 larvae at 20°C or after aHS. (*D-I*) Wide-field images of GFP::LGG-2 in the epidermis of control and *pdzd-8(syb3082)* mutant L4 animals at 20°C (*D*, *G*) or 1h (*E*, *H*) and 3h after aHS (*F*, *I*). (*J-K*) Quantification of the GFP::LGG-2 (*J*) and GFP::LGG-1 (*K*) -positive structures in control and *pdzd-8(syb3082)* at 20°C. One representative of 3 independent experiments is shown. Bars indicate the median, the 25^th^ and 75^th^ percentiles, n animals = 6, 9, 11, 12. Wilcoxon test *p<0.05, NS non-significant. (*L*) Quantification and frequency of GFP::LGG-2 structures with area> 1μm^2^ at 3h post-aHS. Animals n = 33 (ctrl), 44 (*pdzd-8*). LGG-2 puncta n = 165 (ctrl), 531 (*pdzd-8*). Bars indicate the median and the 25th–75th percentiles. Two-tailed Student-t test *** P<0.001. (*M*-*Q*) Confocal images of GFP::LGG-2 autophagosomes and NUC-1::mCherry lysosomes in the epidermis of control (*M*) and *pdzd-8(syb3082)* mutant (*N*), 3h post aHS. Insets are a higher magnification view of the dotted square and show enlarged structures in *pdzd-8(syb3082)* animals. Quantification of the number (*O*) and size (*P*) of lysosomes. Animals n = 6 (ctrl), 8 (*pdzd-8*). NS not significant, ***p<0.01, Wilcoxon test. The colocalization between GFP::LGG-2 and NUC-1::mCherry (*Q*) was calculated with Manders’ M1 coefficient (M1C) from three independent experiments. Bars indicate median, 25^th^ and 75^th^ percentiles, n= 9, 14. * p<0.05, Mann-Whitney U test. (*R*-*W*) Confocal images of GFP::LGG-2 and NUC-1::mCherry 3h post aHS in the epidermis of control (*R*, *U*) and *pdzd-8 (syb3082)* (*S*, *V*) animals submitted to *lgg-1* RNAi (*R*, *S*) or *atg-3* RNAi (*U*, *V*). Manders’ M1 coefficient indicates that the colocalization between lysosome (NUC-1::mCherry, red) and autophagosome (GFP::LGG-2, green) is reduced when autophagy is blocked. Bars indicate median, 25^th^ and 75^th^ percentiles. n=30, 31, 32, 30 (*T*) n= 9, 11, 11, 14 (*W*), Mann-Whitney U test *p<0.05, **p<0.01, ***p<0.001, ***p<0.0001, NS non significative. Scale bar is 10 μm. See also SI-Figure S2, SI-Figure S3.

For the remainder of this study, we focused on the epidermis to analyze the function of PDZD-8 during heat stress-induced autophagy (28). We have previously shown that one hour after an acute heat stress (aHS, one hour at 37°C), a massive transitory autophagy flux was triggered in the epidermis of early fourth-stage larvae (eL4) (28). Because aHS induces ER-phagy (23), we first assessed whether PDZD-8 localization in the epidermis was modified upon aHS administration. Despite a slight decrease in intensity, no substantial modification was observed 1-3 hours after aHS (Fig. 2*A-C*)

Next, we quantified the basal level of autophagy in the epidermis of *pdzd-8(syb3082)* null L4 larvae using as autophagosome markers either GFP::LGG-1 (29) or GFP::LGG-2 (30) and compared it with the control strains. No significant difference was detected for the number of GFP::LGG-1-positive structures (Fig. 2*K* and *SI Appendix*, FigS3*F* *and* *I*). Still, the number of GFP::LGG-2-positive autophagosomes was significantly increased in the *pdzd-8* mutant (Fig. 2*D*, *G*, *J*). One and 3 hours post aHS, the number and the aspect of GFP::LGG-1 puncta did not present a significant difference with the control, indicating that LGG-1 positive autophagosomes are not strongly affected by PDZD-8 depletion (*SI Appendix* Fig. S3*F-L*). On the other hand, abnormally enlarged LGG-2 structures were observed 3 hours after aHS in *pdzd-8* mutant (Fig. 2*E, F, H-L*), supporting a defect in the autophagy flux specific to LGG-2-positive autophagosomes.

The role of LGG-2 in the late stages of autophagosome biogenesis or maturation (9) prompted us to investigate the interaction between the autophagosome and lysosome, using NUC-1::mCherry, which labels the lysosomal lumen (Fig. 2*M-Q*). Three hours after aHS, enlarged lysosomes were present in *pdzd-8* mutant (Fig. 2*M-P*) and accompanied by an increase in colocalization with GFP::LGG-2 (Fig. 2*Q*). An RNAi approach was then performed to check that the abnormal LGG-2 pattern in *pdzd-8* mutant was linked to autophagy. Both LGG-1 and ATG-3 depletion resulted in a substantial decrease of enlarged LGG-2 positive structures in *pdzd-8* mutant, as well as the colocalization with NUC-1 (Fig. 2R-W and *SI Appendix* Fig.S3*M-N*). Altogether, these data suggested that LGG-2 positive autophagosome and autolysosome accumulate in the absence of PDZD-8.

### PDZD-8 promotes the maturation of autolysosomes and the degradation of autophagy cargoes

Time-lapse video-microscopy of GPF::LGG-2 and NUC-1::mCherry was performed after aHS to assess the dynamics of formation of autolysosome in *pdzd-8* mutant (Fig. 3*A* and *SI Appendix* movie 1). When LGG-2-positive vesicles reached the lysosomes, they first clustered around them and only later fused with them, forming enlarged autolysosomes that were positive for both markers. The GFP::LGG-2 fluorescence then progressively diminished, as well as the size of the lysosomes. The identity of LGG-2–positive structures as autophagosomes, NUC-1–positive structures as lysosomes, and double-labelled structures as autolysosomes was confirmed by correlative light and electron microscopy, demonstrating that autolysosome formation is defective in *pdzd-8* mutants. (Fig. 3*B*).

**Figure 3:**
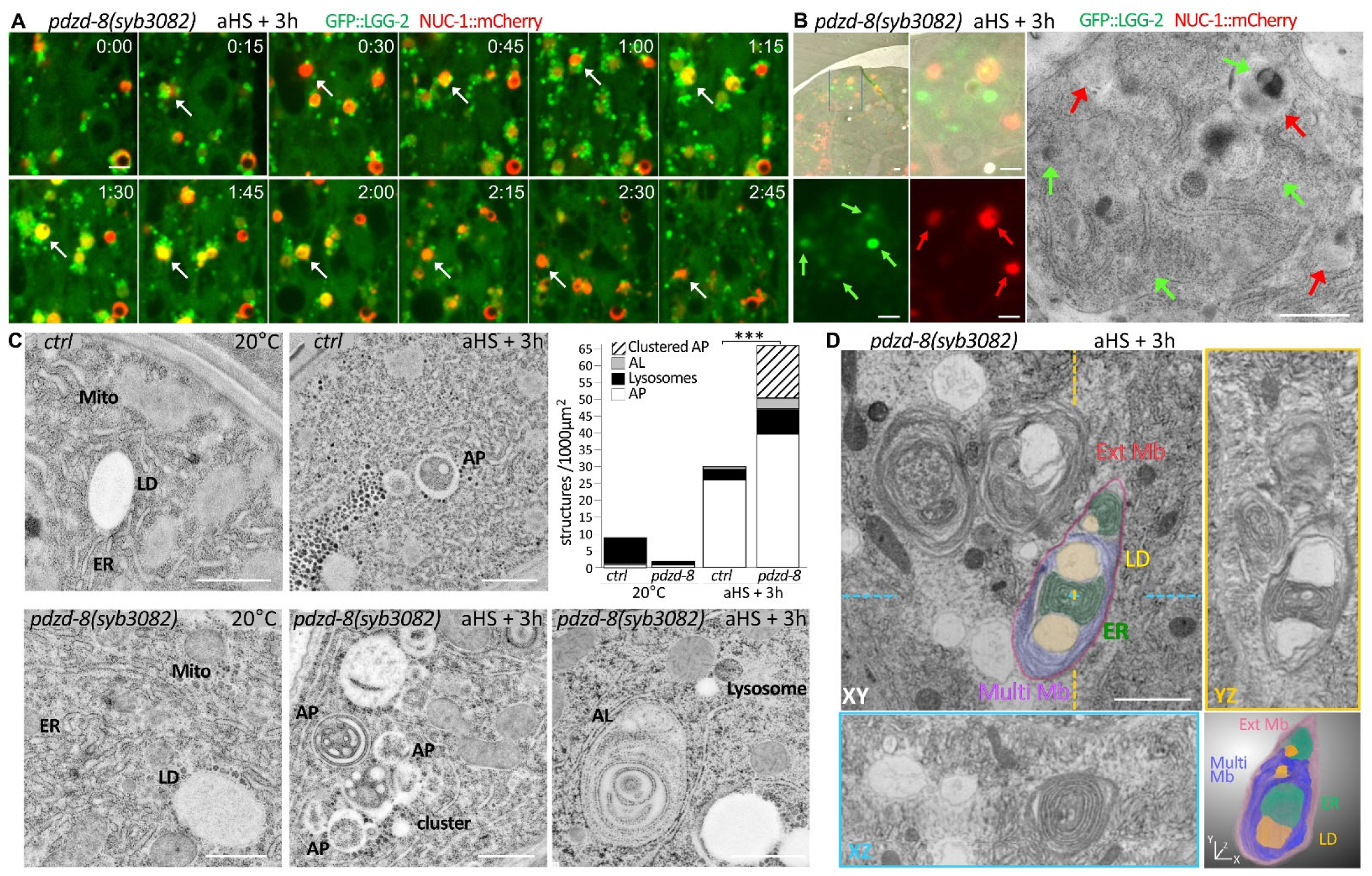
PDZD-8 promotes the formation of the autolysosome and the degradation of autophagic cargoes. (*A*) Time-lapse confocal live imaging of GFP::LGG-2 and NUC-1::mCherry in the epidermis of *pdzd-8(syb3082)* mutant 3h post aHS. The white arrow points to a single lysosome, around which autophagosomes cluster and later fuse, resulting in an enlarged autolysosome (yellow) that remains for an extended period. The GFP::LGG-2 fluorescence finally disappears, and the size of the autolysosome decreases, consistent with the completion of autophagic cargo degradation. See also SI video1. (*B*) Correlative light and electron microscopy (CLEM) analysis of *pdzd-8(syb3082)* mutant epidermis 3h post aHS. Clusters of autophagosome (green arrows) and lysosome (red arrows) of different sizes, whose heterogeneous contents are visible in the magnified corresponding electron micrograph. Image representative of 7 sections from 3 animals. (*C*) Transmission electron micrographs (TEM) of the epidermis of control and *pdzd-8(syb3082)* mutant animals at 20°C or 3h post aHS. In the *pdzd-8* mutant, autophagosomes (AP) with various cargoes often accumulate, forming clusters, and enlarged autolysosomes (AL) with heterogeneous content are present. Quantification of these structures confirms a significant enrichment. n animals= x, n sections= y. Fisher’s exact test. ***p<0.001. (*D*) Focused ion beam–scanning electron microscopy (FIB-SEM) imaging of autolysosome clusters in the epidermis of *pdzd-8(syb3082)* (resolution 10nm). The XY, XZ (blue), and YZ (yellow) plans reveal the presence of packed ER, lipid droplets (LD), and membranous whorls (Multi Mb) inside the autolysosome, whose external membrane (Ext Mb) is highlighted in red. A 3D rendering illustrates the intrication of the different structures within the autolysosome. See also SI Fig.S4, SI movies2 and 3. Scale bar is 1µm.

A quantitative transmission electron microscopy approach of the epidermis of control and *pdzd-8* mutant was also used to document the contents of the autophagosomes and the autolysosomes (Fig. 3*C*). This analysis revealed that the cargoes already described in aHS-induced autophagy (23) are detected in the autophagosomes that are clustered with lysosomes but also in enlarged autolysosomes of *pdzd-8* mutant. This data confirmed that PDZD-8 depletion affects the fusion of autophagosomes with lysosomes and their subsequent degradation, but not the formation of autophagosomes or the sequestration of cargoes.

Finally, a FIB-SEM was performed to obtain a high-resolution 3D reconstruction of the autolysosome shape and content in the *pdzd-*8 mutant (Fig. 3D and *SI Appendix* Fig. S4, movies 2 and 3). This analysis demonstrated the dense packing of multiple and varied compartments within the autolysosome, including multilamellar ER, vesicles with undefined cytoplasmic contents, membrane whorls, and lipid droplets. These observations suggested that aHS induces ER-phagy and lipophagy and that this process can still be initiated in the absence of PDZD-8. However, despite lysosomal delivery, lipid droplets appear to remain largely intact within lysosomes, indicating a defect at a later degradative step.

Altogether, these results support a role for PDZD-8 in the fusion of the autophagosome with the lysosome and in the degradation of autophagic cargoes.

### Autolysosome formation and maturation rely on PDZD-8 interaction with LGG-2

To determine whether the autophagy-related function of PDZD-8 depends on its ability to interact with LGG-2, we generated a CRISPR-Cas9-engineered EH**A**EE**A** mutant of the LIR motif, called *pdzd-8(LIR)*. We also used *pdzd-8(ΔSMP)* mutant, an in-frame deletion of the SMP domain, to test whether the lipid transfer capacity was required for PDZD-8 autophagy functions (19). We compared GFP::LGG-2 and NUC-1::mCherry pattern at 20°C and 3 hours after aHS in *pdzd-8(LIR)*, *pdzd-8(syb3082)* and *pdzd-8(ΔSMP)* mutants (Fig. 4*A* and *B*). Both the *pdzd-8(LIR)* and *pdzd-8(ΔSMP)* mutants led to the formation of large autolysosomes at levels comparable to those observed in the *pdzd-8* null mutant, suggesting that the two domains are necessary for autophagy-related functions of PDZD-8. Immunofluorescence showed that the localization and amount of PDZD-8(LIR) protein were similar to those of the wild-type protein, indicating that PDZD-8 interaction with LGG-2 is essential for its function in autophagy (*SI Appendix* Fig. S5*A-C*). Moreover, the deletion of the LIR motif significantly reduces the colocalization between LGG-2 and PDZD-8, confirming that the LIR is required for PDZD-8 interaction with LGG-2. (*SI Appendix*, Fig. S5*F-H*). However, the amount of PDZD-8(ΔSMP) was significantly diminished (*SI Appendix* Fig. S5*D-E’*), indicating that the SMP domain is required for PDZD-8 localization at the ER, making it difficult to conclude on the autophagy function of the SMP domain *per se*.

**Figure 4:**
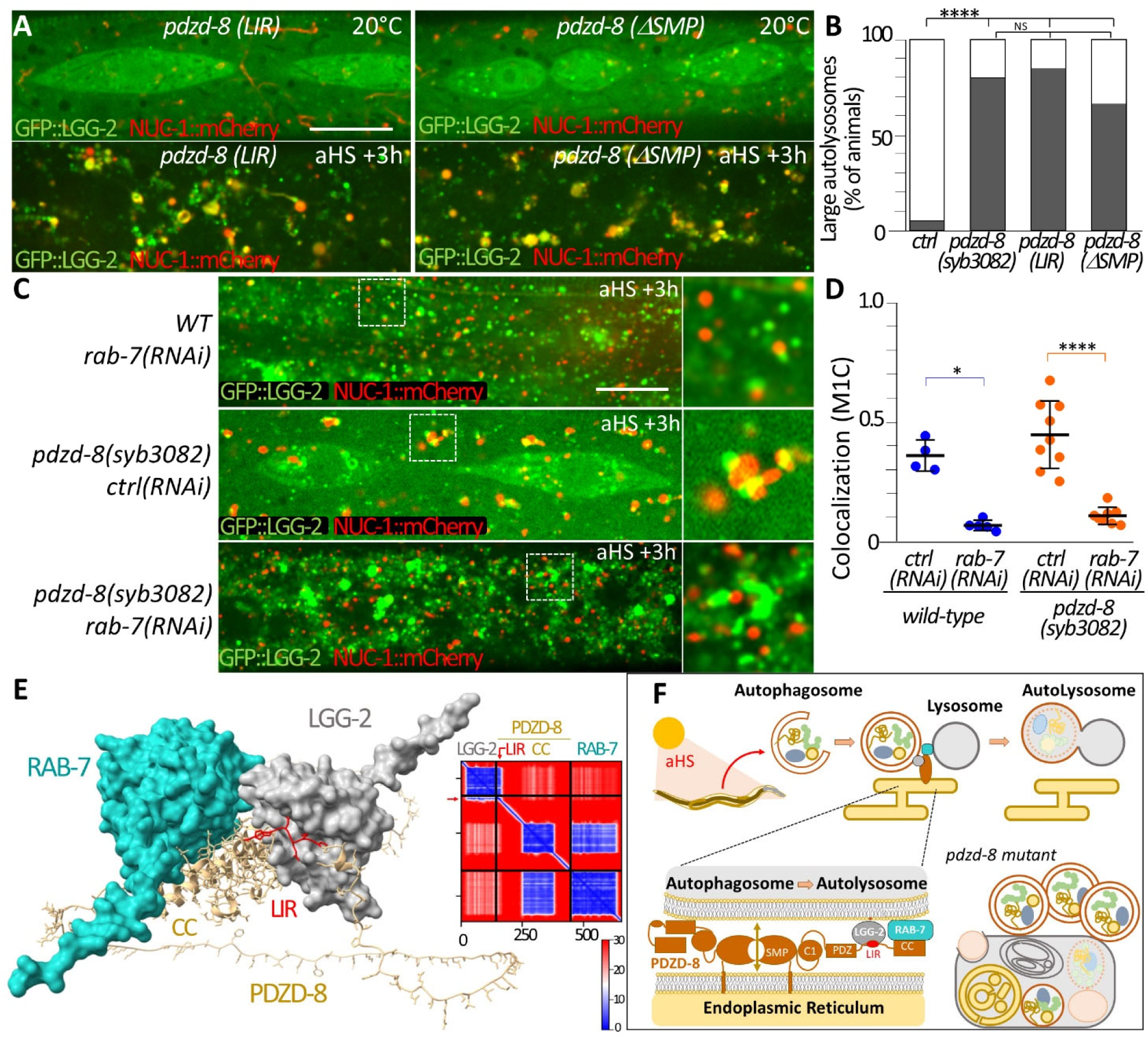
Autolysosome formation and maturation rely on PDZD-8 interaction with LGG-2. (*A*, *B*) Spinning confocal images of GFP::LGG-2 and NUC-1::mCherry in the epidermis of *pdzd-8 (syb922(LIR)*) and *pdzd-8(syb2977)(ΔSMP*) mutants at 20°C or 3h post aHS (*A*) and quantification (*B*). The frequency of worms harboring large autolysosomes in *LIR* and *ΔSMP* mutants is similar to that *of* the null mutant *syb3082*. n animals = 19, 10, 13, 9. Multiple Pearson’s χ²test with Yates’ continuity correction and Bonferroni correction, ****p<0.0001, NS: non-significant. (*C*, *D*) Confocal images of GFP::LGG-2 and NUC-1::mCherry in the epidermis of control and *pdzd-8* null mutant 3h post aHS (*C*) submitted to *rab-7* RNAi and quantification (*D*). Manders’ M1 coefficient (M1C) indicates that the colocalization between lysosome and autophagosome in *pdzd-8* is reduced when RAB-7 is depleted. Bars indicate median, 25^th^ and 75^th^ percentiles. n=4, 5, 9, 8 Mann-Whitney U test *p<0.05, ***p<0.0001. See also SI-Figure S5. (*E*) AlphaFold 2.3 model of the LGG-2/PDZD-8/RAB-7 tripartite complex and predicted aligned error (PAE) matrix. (*F*) A proposed model schematic for PDZD-8 role in aHS-induced autophagy. PDZD-8 is recruited to ER-autophagosome and/or ER-autolysosome contact sites via the LC3/LGG-2 autophagy protein and Rab7 respectively. These contact sites may facilitate lipid transfers (double brown arrow) between ER and autophagosome/lysosome to permit the fusion to form autolysosome. The loss of PDZD-8 at these contact sites resulted in a transient autophagy blockage due to a delay within the process of fusion between autophagosome and lysosome leading to the formation of a cluster of intermingled autophagosome and lysosome. Scale bar: is 10 μm.

Finally, we tested whether the function of the GTPase RAB-7 was involved in the autophagy-related functions of PDZD-8. RAB-7 is well characterized for its role in autophagosome-lysosome fusion (9, 31, 32) and is a direct interactor of the coiled-coil domain of PDZD-8 (13–15). Moreover, an AF2 model with a high confidence score supported the possibility of a tripartite complex with PDZD-8, LGG-2, and RAB-7 (Fig.4*E*). RAB-7 was depleted by RNAi in the *pdzd-8* mutant at 20°C and after aHS, and GFP::LGG-2 and NUC-1::mCherry were analysed (Fig. 4*C* *and* *D*). The inactivation of *rab-7* blocked the autophagy flux, leading to a significant increase in LGG-2 autophagosomes and almost no co-localization with NUC-1, both in the control and in *pdzd-8* null mutant. This result indicates that *rab-7* inactivation suppresses the *pdzd-8* mutant phenotype, supporting a PDZD-8 function downstream of RAB-7. In summary, our data support a novel function of PDZD-8 during autophagy, promoting the fusion of autophagosomes with autolysosomes and the degradation of autophagy cargoes.

## Discussion

PDZD-8 regulates organelle homeostasis by organizing membranes at the ER-endosome/lysosome (12, 13, 15, 18, 19, 22) and ER-mitochondria contact sites (16, 17). This study identifies PDZD-8 as a new interactor of LGG-2/LC3, which modulates autophagosome/autolysosome maturation during heat stress-induced autophagy.

Our EM analyses of autophagic cargoes in the *pdzd-8* mutant confirmed that multi-lamellar ER stacks are sent for degradation by ER-phagy (23). However, the accumulation of numerous lipid droplets is the first evidence that aHS induces lipophagy. The nature of autophagy has yet to be determined, but the presence of not fully engulfed lipid droplets in large invaginations of the autolysosome border (*SI Appendix* Fig.S4) is suggestive of microlipophagy. This is reminiscent of the phenotype observed in brain-specific PDZD8 knockout mice, where undegraded lipid droplets accumulated in proximity to lysosomes (21). This defect was interpreted as an impairment of the lipid droplet and lysosome fusion, consequent to defective endo-lysosomal maturation. This is a supplementary indication that aHS induces several types of selective autophagy concomitantly to cope with a harsh stress affecting multiple cell structures.

The PDZD-8 protein is weakly expressed in *C. elegans* and is predominantly present in the nervous system. Similar observations in flies (22) and mice (21) support an evolutionarily conserved role, in agreement with genetic evidence in humans associating variants of PDZD8 with neurodevelopmental and cognitive impairments (33–35). Interestingly, LGG-2, but not LGG-1, is also enriched in neurons in *C. elegans* (30).

After acute heat stress, two main phenotypes were observed in *pdzd-8* mutants. First, autophagosomes frequently formed clusters around lysosomes, supporting a delayed fusion between these compartments. Second, the accumulation of cargoes and persistence of GFP::LGG-2 fluorescence inside enlarged autolysosomes revealed that their degradative capacity was affected. Our data indicate that PDZD8 does not control the initiation of autophagy but accelerates late autophagic steps through its role at ER membrane contact sites. The formation of enlarged autolysosomes was suppressed by *rab-7* RNAi, suggesting that the *pdzd-8* phenotype manifests downstream of autophagosome–lysosome tethering.

We cannot exclude that the defect in the degradation of cargoes is independent of the binding of PDZD-8 to LGG-2. A recent study demonstrated that PDZD8 can transfer large amounts of lipid from the ER to lysosomes to accommodate its membrane expansion under osmotic stress (20). Moreover, a recent study in the fly *D. melanogaster* proposed that PDZD8-mediated ER-lysosome contact sites promote efficient lysosomal activity and enhance autophagic flux (22). In this regard, the *pdzd-8* mutant phenotypes observed after aHS probably resulted from the combination of autophagosomal and lysosomal dysfunctions. In the absence of PDZD-8, the rapid and massive delivery of autophagic cargoes overwhelms the lysosome’s degradative capacity. However, the LGG-2-specific LIR motif is essential for PDZD-8 function in autophagy, indicating that PDZD8 directly acts at ER-autophagosome contact sites, and not only at ER-lysosome interfaces. PDZD8, in association with LGG-2, could organize a new type of contact zone between the ER and autophagosomes, or potentially very recently formed autolysosomes that still retain LGG-2 on their surface. PDZD-8 tethering with autophagosome is specific for LGG-2/LC3 but not for LGG-1/GABARAP, further confirming the dedicated function of LGG-2 in late autophagosome maturation in *C. elegans* (16). In particular, LGG-2 was shown to interact with the HOPS subunit VPS-39, facilitating the RAB-7-dependent tethering of lysosomes during a developmental mitophagy process. Our study is a new example of LGG-1 and LGG-2 distinct functions in autophagy, in agreement with the differential roles of mammalian LC3 and GABARAP families in autophagosome biogenesis and maturation (36, 37)

ER-autophagosome interactions are well described for the initiation and elongation of the isolation membrane, which gives rise to the autophagosome. The late-stage-specific functions of the PDZD-8-LGG-2 tether imply tight regulation, either through the reorganization of pre-existing ER-autophagosome contact sites or the de novo formation of new ones at later steps of autophagy. Altogether, this supports a model in which autophagosomes remain closely associated with the endoplasmic reticulum throughout their maturation, indicating that sustained ER interactions are required for the progression of autophagy.

What is the function of PDZD-8 at the ER-autophagosome contact site? The size and the presence of multiple binding domains in PDZD-8 support a structural function in tethering the ER with the autophagosome and the lysosome. However, the tissue-specific and low level of expression, as well as the phenotype in basal conditions, indicate that tethering between autophagosomes and lysosomes is not the primary function of PDZD-8.

PDZD-8 is well characterized to transport lipids between ER and endo-lysosomal membranes, and in *C. elegans*, it mediates the removal of PI(4,5)P₂ from recycling endosomes (19). Similarly, PDZD-8 could optimize the lipid composition and membrane properties of the autophagosome, thereby conditioning its competence for efficient fusion. Indeed, the fusion between autophagosomes and lysosomes is dependent upon the lipid composition of both autophagosomal and lysosomal membranes (10, 38). Alternatively, the lipid composition of the autolysosome membrane, which is derived from lysosomes, endosomes, and autophagosomes, could influence the subsequent final maturation of this hybrid organelle. It may affect the recruitment of autophagosomal components recycling (ACR), which include SNX4, SNX5, and SNX17 that have specific binding capacities for phosphoinositides employed for autolysosomal membrane recruitment (39). Interestingly, the blockage of ACR inhibits the autophagy flux. Another possibility is that PDZD-8 regulates autophagosome and autolysosome function by modulating their lipid composition, thereby directly impacting autolysosome maturation and degradative capacity. In yeast, PI(3,5)P₂ promotes the stabilization and assembly of v-ATPase subunits on lysosomal membranes, leading to enhanced proton pumping, maintenance of an acidic luminal pH, and efficient activation of hydrolytic enzymes (40). By analogy, PDZD-8–dependent lipid remodeling could control v-ATPase activity at late autophagy stages and thus tune autolysosomal proteolysis.

## Materials and Methods

Detailed procedures and protocols used in this study are described in the *SI Appendix*.

### *C. elegans* strains and growth conditions

*C. elegans* strains were maintained on NGM agar plates seeded with *E. coli* OP50 at 15 °C, 20°C, or 25 °C using standard methods (51). N2 Bristol was used as the wild-type reference strain. Synchronized populations were obtained by timed egg laying. Heat stress experiments were performed on synchronized early L4 larvae by incubation at 37 °C for 1 h, followed by recovery at 20 °C. All strains used in this study, including CRISPR-engineered alleles, fluorescent reporters, genetic crosses, and genotyping strategies, are listed in the *SI Appendix*.

### Yeast two-hybrid screening

Protein–protein interactions were identified using a yeast two-hybrid (Y2H) screen with full-length LGG-2 as bait against a mixed-stage *C. elegans* cDNA library (Hybrigenics Services). Positive interactions were selected under stringent conditions, assigned confidence scores, and validated by independent one-to-one Y2H assays.

### Structural modeling

Structural predictions were generated using AlphaFold2 to model the interaction between LGG-2 and the C-terminal region of PDZD-8 containing the LC3-interacting region (LIR). Structural visualization and analysis were performed using ChimeraX.

### Antibody production and immunolocalization

Polyclonal antibodies against PDZD-8 were generated using immunogenic peptides selected based on AlphaFold predictions (Eurogentech service). Immunolocalization experiments were performed on early embryos using freeze-fracture and methanol fixation, followed by staining with primary and fluorescent secondary antibodies.

### Light and confocal microscopy

Fluorescence imaging of live and fixed samples was performed using widefield epifluorescence, confocal microscopy and confocal spinning-disk microscopy. Live animals were immobilized either on agarose pads or in microfluidic devices, depending on the experiment. Imaging parameters were adjusted according to experimental conditions. Microscope configurations and acquisition settings are detailed in the Supporting Information.

### Electron microscopy and correlative imaging

Transmission electron microscopy (TEM), correlative light and electron microscopy (CLEM), and focused ion beam–scanning electron microscopy (FIB–SEM) were performed following high-pressure freezing and freeze-substitution. Three-dimensional reconstructions were generated from FIB–SEM datasets. Complete sample preparation, imaging, and image-processing workflows are provided in the Supporting Information.

### RNA interference

RNA interference experiments were performed using bacterial feeding with HT115 (DE3) strains that express double-stranded RNA targeting specific genes. Synchronized F1 progeny were analysed at the early L4 stage.

### Phenotypic analysis

Phenotypic analyses, including body size measurements, developmental assessments, and lifespan assays, were performed on synchronized animals.

### Image analysis and statistics

Image processing, segmentation, and quantification were performed using ImageJ/Fiji. Colocalization analyses were conducted using Manders’ coefficients. Statistical analyses were performed using GraphPad Prism or R. Full image analysis pipelines and statistical methods are described in the Supporting Information.

## Author Contributions

RL acquired funding and led project administration. EC and RL conceived the project and designed experiments. AP, VM, EC, CLe and CLa performed the experiments. AlphaFold analysis was conducted by AP, EC and DJZ. TEM, CLEM and FIB-SEM experiments and image processes were conducted by CLa with the help of CB and BR for FIB-SEM. AP, VM, EC, CLa, CLe, and RL analysed and interpreted the data. EC, AP and RL prepared the first draft of the manuscript with input and approval from all authors.

## Acknowledgments

We thank Yasunori Saheki (Nanyang Technological University, Singapore, and Kumamoto University, Japan) for providing strains PHX2977 and SAH607. We thank the Caenorhabditis Genetics Center, which is funded by the National Institutes for Health Center for Research Program, for providing some of the *C. elegans* strains used in this study. We thank the BIOI2 platform for making the ColabFold pipeline easily accessible at the I2BC. We thank Nicolas Regnard and Siam Siraji for technical help with the construction and initial characterization of strains. We are grateful to the members of the Legouis lab (Marie-Hélène Cuif, Magali Prigent, Laurent Kuras, and Claudia Serot) for providing constructive and helpful feedback throughout this work. This work benefited from the core facilities of Imagerie-Gif (http://www.i2bc.paris-saclay.fr), a member of Infrastructures en Biologie Santé et Agronomie (http://www.ibisa.net), supported by France BioImaging (ANR-10-INBS-04-01) and the Labex Saclay Plant Science (ANR-11-IDEX-0003-02). This work was supported by the Agence National de la Recherche (grant numbers ANR-23-CE13-0013-01) and AP by a Ph.D. internship from CNRS

## Supporting Information Text

### Results

#### *pdzd-8* locus codes for 11 putative isoforms

Alternative transcription initiation (a1 and a2) and termination (h1 and h2) sites, along with alternative splicing, lead to 13 different transcripts encoding 11 protein isoforms ranging from 1264 to 1365 amino acids. By aligning the protein sequences of the different isoforms, we identified differences between them. Subsequently, analysis of the corresponding transcripts identified the molecular processes that underpin the generation of the diverse transcripts, which in turn give rise to the different isoforms. The K isoform is the longest, comprising 1,365 amino acids. This isoform serves as a reference point for illustrating the observed changes in all the other pDZD8 isoforms. The F isoform lacks 50 amino acids at position 425 due to the absence of exon 7 in its transcript. Furthermore, exon 11 is also truncated. The 3’ splice site of the intron situated before exon 11 is shifted to 3’, which eliminates a sequence of exon 11 that follows, namely, six nucleotides, corresponding to the FQ dipeptide. Furthermore, the intron splice site located immediately upstream of exon 18 has undergone a 5’ shift. This results in the addition of two amino acids and a phase shift, introducing an early stop codon and truncating the C-terminal end, yielding the sequence RRLKQ. Transcripts a1 and a2 encode the same A isoform. Indeed, the sole distinction between these two isoforms is the 5’ untranslated region (UTR). The a2 transcript is seven nucleotides longer. This suggests the existence of at least two distinct transcription start points. The two transcripts, designated h1 and h2, differ by two nucleotides at their 3’UTR. However, both transcripts encode the same H isoform, which has a truncated C terminus and the following amino acid sequence: RRLKQ. Isoform I also exhibits a truncated C-terminus, terminating in RRLKQ. Additionally, the transcript that forms the I isoform exhibits a slightly altered exon 17 sequence. Two 3’ intron splice sites were identified for the intron preceding exon 17. Transcripts of isoform I utilise the most 3’ located splice site, resulting in exon 17 having a truncated ACATTTCAG sequence and lacking the ISE amino acid sequence. The remaining isoforms are produced from transcripts that combine the indicated alternative splicing events. Isoform A is devoid of both the FQ and ISE motifs. Isoform B is deficient in the ISE motif. Isoform D is lacking both the FQ and ISE sequences and is devoid of exon 7. Isoform E is deficient in the FQ motif and has a truncated C-terminal ending with RRLKQ. Isoform G is lacking in the ISE sequence and does not contain exon 7. Isoform J is deficient in both the FQ and ISE motifs and has a truncated C-terminal ending with RRLKQ.

### Extended materials and methods

#### Strains, growth conditions, genetic crosses, and genome editing

The *C. elegans* strains were cultivated at 25°C, 20°C, or 15°C on NGM agar plates seeded with the E. coli strain OP50, following standard protocols (1). The *C. elegans* strain N2 Bristol was used as the control (wild-type) in this work. Synchronized *Caenorhabditis elegans* populations were obtained by allowing gravid hermaphrodites to lay eggs for 1.5 h at 20 °C. Synchronized early L4-stage animals, approximately 44 h after egg laying, were subjected to heat stress. Heat stress was induced by incubating worms at 37 °C for 1 h, followed by a recovery period of 1h to 3h at 20 °C.

We employed the following *C. elegans* genetic strains. RD475*[P_lgg-2_::gfp::lgg-2]IV; [P_nuc-1_::mCherry::nuc-1]* to visualise both autophagosomes and lysosomes and their respective locations *in vivo*. *ocfIs2[pie-1p:mCherry::sp12::pie-1 3’UTR + unc-119(+)]* to visualise the endoplasmic reticulum within the embryo cells. PHX2977:*tex-2(syb670)II;esyt-2(syb709)III;pdzd-8(syb2977)IV;tmem-24(tm10626)X;zuIs45[nmy-2::NMY-2::GFP+unc-119(+)]V* and SAH607: *syb4099[PDZD-8::mNG])IV*. Both strains are a gift from Yasunori Saheki’s lab. PHX2977 is a quadruple mutant of the only 4 *C. elegans* genes that encode proteins with an SMP domain. It combines null alleles of *tex-2*, *esyt-2*, and *tmem-24*, and a mutant allele of pdzd-8 that selectively deletes the coding region for the SMP domain. This mutant allele therefore encodes a PDZD-8 protein that lacks the SMP domain and therefore lacks PDZD-8 lipid transfer activity. We crossed PHX2977 with RD475 to generate RD504: *pdzd-8(syb2977)IV,Is[GFP::LGG-2]IV;Is[NUC-1::mCherry]*. Animals were selected by PCR genotyping to identify those that have recovered a wild-type phenotype for *tex-2*, *esyt-2*, and *tmem-24* but retain the SMP deletion in the *pdzd-8* gene. SAH607 expressed a mNeonGreen-tagged PDZD-8 at the *pdzd-8* locus. We out-crossed SAH607 with N2 to get strain RD358: *syb4099*[PDZD-8::mNG])IV.

SunyBiotech Corporation (Fuzhou, China) generated a CRISPR-Cas9-engineered strain used in this study. These strains were outcrossed at least 3 times with N2 before being used for further studies. PHX3082: *pdzd-8 (syb3082)*, PHX3106: *syb3106[PDZD-8::GFP])IV*. PHX3106 could be used to visualise the endogenous expression pattern of PDZD-8. However, no or very weak GFP signal was detected in this strain, precluding its usefulness. PHX-9229: *pdzd-8(syb9229)[F1065A,I1068A]IV.* PHX9229 has two point mutations in the *pdzd-8* gene. These point mutations affect the LIR motif of the PDZD-8 protein. These mutations are associated with a restriction polymorphism (cleavage site for the restriction enzyme PaeI), which permits the monitoring of the segregation of this mutant allele during genetic crosses. PHX3098: *ret-1(syb3098)[pan RET-1::wrmScarlet]V* permits visualizing the endoplasmic reticulum in most of the *C. elegans* cells.

We crossed PHX3082 with RD224 Is*[NUC-1::mCherry]*; Is*[lgg-2::gfp::lgg-2]IV* to create RD451: *pdzd-8(syb3082)IV;*Is*[lgg-2::gfp::lgg-2]IV; [nuc-1::mCherry::nuc-1].* RD224 and RD451 were outcrossed twice with N2 to produce RD475 and RD476, respectively. We crossed PHX3082 with *Is[patg-18::ATG-18::GFP]* to get RD455: *pdzd-8 (syb3082)IV;Is[patg-18::ATG-18::GFP]*. We crossed PHX3082 with DA2123: *adIs2122[lgg-1p::GFP::lgg-1;rol-6(su1006)]* to get RD457: *pdzd-8 (syb3082)IV; adIs2122[gfp::lgg-1; rol-6(su1006)]*. We crossed PHX-9229 with RD475 to get RD503:*pdzd-8(syb9229)[F1065A,I1068A]IV,*Is*[lgg-2::gfp::lgg-2]IV;[nuc-1::mCherry::nuc-1]*. We crossed *Is[GFP::LGG-2]* with PHX3082 to get RD460: *pdzd-8 (syb3082)IV*, *Is[GFP::LGG-2]V*. We crossed PHX3082 with PHX3098: *ret-1(syb3098)[pan RET-1::wrmScarlet]V* to get RD458: *pdzd-8 (syb3082)IV*; *ret-1(syb3098)[pan RET-1::wrmScarlet]V.* We crossed *PHX3098* with SAH607: *syb4099[PDZD-8::mNG])IV* to get RD474: *pdzd-8(syb4099[PDZD-8::mNG])IV;ret-1(syb3098)[pan RET-1::wrmScarlet]V*.

Details on the protocols used for genotyping the different alleles reported in this work are available upon request (primer sequences, PCR conditions, amplicon sizes from putative null mutants and corresponding wild types).

#### Yeast two-hybrid (Y2H) screening and assays

Protein-protein interaction analysis was performed using the yeast two-hybrid (Y2H) system in collaboration with Hybrigenics Services (Paris, France). The Y2H screening was conducted to identify interacting partners of LGG-2, the bait. A full-length cDNA of lgg-2 was cloned into the pB27 vector (Hybrigenics) downstream of the LexA DNA-binding domain (BD) to create the bait construct. This construct was transformed into the yeast strain L40ΔGal4 (MATa, trp1, leu2, his3, LYS2::(lexAop)4-HIS3, URA3::(lexAop)8-lacZ). For the prey library, a C. elegans mixed-stage random-primed cDNA library fused to the Gal4 activation domain (AD) in the pP7 vector was used. Yeast cells containing the bait construct were mated with yeast cells carrying the prey library. Interaction screening was performed under selective conditions lacking leucine, tryptophan, and histidine, supplemented with 3-amino-1,2,4-triazole (3-AT) to inhibit any background HIS3 expression. Positive colonies were identified by their ability to grow on selective media containing 100 mM 3-AT. Positive clones were isolated, and prey plasmids were recovered and sequenced. The sequences were analyzed with Hybrigenics’ bioinformatics pipeline to identify putative interacting proteins. The strength of interactions was classified into four confidence levels (A-D; A having the highest confidence in binding) based on Hybrigenics algorithms. For confirmation, selected interactions were further validated by re-cloning and performing independent one-to-one Y2H assays with co-transformation of bait and prey constructs in the L40ΔGal4 strain.

#### AlphaFold Structural Prediction and Protein Interaction Analysis

To investigate the potential interaction between LGG-2 and PDZD-8, structural predictions were performed using AlphaFold2 (2). We employed the I2BC home-made Alphafold server (ColabFold v1.5.2 using Alphafold 2.3). The full-length amino acid sequences of LGG-2/LC3 proteins were obtained from the UniProt database. We specifically focus on the last 400 amino acids of PDZD-8 (isoform K), which contain the SID region and the LC3-interacting region (LIR) motif.

For structural analysis, the predicted interaction between the LIR domain of PDZD-8 and LGG-2 was imported into PyMOL, and the binding groove was analyzed by visualizing potential hydrogen bonds, hydrophobic contacts, and other stabilizing interactions. Key residues at the interface were highlighted. Analysis of the structural models was conducted by evaluating confidence scores provided by AlphaFold for each residue within the predicted structure. High-confidence regions (pLDDT > 80) were used to interpret the interaction with confidence. All figures showing the predicted interaction between LGG-2 and PDZD-8 SID were generated in PyMOL, and the images were annotated to indicate the LIR domain and the critical residues involved in the interaction. Additionally, to validate the functional relevance of the interaction, conservation of the LIR motif in PDZD-8 was assessed by comparing the predicted LIR domain from PDZD-8 proteins from other nematode species. Sequence alignment was performed using Clustal Omega.

#### Polyclonal PDZD8 Antibody Production Using Immunogenic Peptides

This PDZD-8 polyclonal antibody production was performed in collaboration with Eurogentec. Using AlphaFold, we identify suitable immunogenic regions at the surface of the PDZD-8 protein, typically 15 amino acids in length. We initiated two immunisation programs, one with peptides NH2-PTKLDPSSDAVEAHS (P1351-S1365) and NH2-SVNVDSNNEEDSESV (S1316-V1330), and the other with a mix of peptides NH_2_-ISNRTDTGTEDGGDD (I646-D660) and NH_2_-KRKNTDASDLNGESI (K696-I710). Rabbits were immunized with the peptide-KLH (Keyhole limpet hemocyanin) conjugate. We purify antibodies from the serum using affinity chromatography with the immunization peptides. We tested the purified serum and found that the batch from the rabbit immunized with NH_2_-ISNRTDTGTEDGGDD and NH_2_-KRKNTDASDLNGESI, purified with the column containing peptide ISNRTDTGTEDGGDD, gave the best result. Purified antibody is stored at −20°C with glycerol.

#### Immunolocalization

To release early embryos, 100 adult hermaphrodites were cut on a poly-L-lysine-coated slide (0.1%). Embryos were prepared for immunofluorescence by freeze-fracture and methanol fixation for 30 minutes at –20 °C, followed by a 40-minute incubation in a solution of 0.5% Tween and 3% BSA in PBS. The embryos were washed twice for 30 minutes in 0.5% Tween-PBS. Primary antibodies (anti-PDZD-8 1/250, anti-GABARAP, and anti-LGG-2, anti-GFP 1/250, anti-mNeonGreen 1/250) were applied overnight at 4 °C. After two washes, secondary antibodies (Alexa 488 1/500 and Alexa 643 1/200) were incubated for 2 hours at room temperature, followed by two additional washes. Finally, embryos were mounted in DABCO and imaged.

#### Light microscopy imaging

Epifluorescent images were acquired using a Zeiss AxioImager M2 microscope, equipped with either a 10×/0.3 NA EC Plan Neofluar or a 100×/1.4 NA Plan Apochromat DIC oil objective. Image acquisition was managed using ZEN software (Zeiss) and an AxioCam506 mono camera. To immobilize the worms, they were placed on 2% agarose pads in 2 µL of M9 buffer containing 40 mM sodium azide. Imaging was restricted to a maximum duration of 15 minutes post-mounting. Images of fixed embryos from immunolocalization experiments were acquired with a Leica confocal microscope. For imaging live worms, we employed a Nikon Inverted Eclipse Ti-E confocal microscope equipped with a spinning disc (CSU-X1-A1, Nipkow Spinning Disk confocal system (Yokogawa) with a 100x PLAN APO oil immersion objective (N.A: 1.40). Image acquisition was controlled by a camera Prime 95B sCMOS (Photometrics) with 11µmx11µm pixel size operated through MetaMorph Software version 7.7 (Molecular Devices). Image acquisition was performed in a stack mode with a 0.5 µm step size. When we imaged the epidermis, we focused on the anterior part of the animal, posterior to the terminal bulb. To analyse the expression pattern of PDZD-8::mNeonGreen, image stacks were acquired along the entire anteroposterior axis of the animal. The acquisition parameters were set as follows: GFP channel – 20% laser (488 nm) power with a 500 ms exposure time, and mCherry channel – 10% laser (561 nm) power with a 500 ms exposure time.

#### Confocal spinning disk microscopy live imaging

Heat-stressed, synchronized early L4 animals were collected from plates after the indicated recovery period using M9 buffer. The suspension was centrifuged for 30 s at 1000g to concentrate the animals. Worms were then loaded into microfluidic chips designed for immobilizing L4-stage worms (Vivoverse, vivoChip-2x L4-YA). Submitted to a continuous flow of M9 buffer, applied at a pressure of 3.5 psi. Then, microfluidic devices were mounted on the microscope stage, and image acquisition was initiated once animals were stabilized within the channels. Acquisition parameters were adjusted depending on the experimental setup.For experiments shown in Fig. 3A, imaging was performed using a 100× objective without a Perfect Focus System (PFS). Fluorescence was detected using a dual-band 525/45 nm filter for 488 nm excitation and a dual-band 607/36 nm filter for 561 nm excitation. The 488 nm laser was set to 10% power, and the 561 nm laser to 15%, with an exposure time of 300 ms per channel. Z-stacks were acquired with a 0.5 µm step size over 15 optical sections. Images were collected every 15 min. For experiments shown in Fig. S3 and the associated time-lapse video, imaging was performed using a 60× objective equipped with a Perfect Focus System. Fluorescence was detected using the same dual-band filters (525/45 nm for 488 nm excitation and 607/36 nm for 561 nm excitation). Both lasers were set to 3% power, with an exposure time of 300 ms. Z-stacks were acquired with a 0.5 µm step size over 15 optical sections, and images were collected every 5 min.

#### Confocal microscopy live imaging

PDZD-8:mNeonGreen fluorescence imaging was performed using a standard confocal microscope (Leica SP8-X) equipped with a white light laser. The mNG fluorescence was excited at 503 nm, with the laser power set to 30%. Images were acquired using an exposure time of 500 ms,

#### Transmission Electron Microscopy

Control and mutant nematodes, either maintained at 20°C or subjected to heat shock, were transferred into 200 µm deep flat carriers pre-coated with 1% phosphatidylcholine in chloroform (Sigma-Aldrich) and immersed in 1µl-2µL of 20% BSA (Sigma-Aldrich, A7030) in M9 buffer. The samples were then cryo-immobilized using the EMPACT-2 high-pressure freezer (Leica Microsystems, Vienna, Austria) following the method described in (3). Cryo-substitution was carried out with an Automated Freeze Substitution System AFS2 (Leica Microsystems, Wetzlar, Germany). The carriers were placed in the AFS2 chamber, pre-set at −90°C, with cryo-substitution buffer (acetone, 1% osmium tetroxide, 0.25% uranyl acetate) in cryogenic tubes. The samples remained at −90°C for 30 hours, after which the temperature was gradually increased to −30°C at a rate of 3°C per hour. Once at −30°C, the samples were maintained for 12 hours before being brought to 0°C. The samples were then removed from the AFS2 and kept on ice at 4°C, followed by four 10-minute washes in anhydrous acetone. Next, the samples were infiltrated with EPON (Agar Scientific, R1165) at room temperature in a series of increasing concentrations: 25% for 4 hours, 50% for 4 hours, 75% overnight, and 100% for 8 hours, followed by another overnight incubation in 100% EPON. The samples were then embedded in fresh EPON and cured at 60°C for 24 hours. Ultrathin sections (80 nm) were cut using an ultramicrotome (Leica Microsystems, EM UC7) and collected on formvar and carbon-coated copper slot grids (LFG, FCF-2010-CU-50). The sections were stained with 2% uranyl acetate for 15 minutes and 0.08 M lead citrate (Sigma-Aldrich, 15326) for 8 minutes. The sections were examined with a JEOL 1400 transmission electron microscope (TEM) at 80 kV, and images were captured using a Gatan SC1000 Orius 11-megapixel CCD camera.

#### CLEM

CLEM experiments were performed as described in (4) with the following modifications. Cryo-substitution was carried out at −90°C in acetone containing 0.01% osmium tetroxide. Dehydration was performed in 95% acetone/5% water supplemented with 0.1% uranyl acetate. The solvent was progressively exchanged for glycol methacrylate (GMA) resin (30%, 70%, and 100%) in 95% ethanol/5% water at −30°C. Resin polymerization was carried out at −30°C using *N, N*-dimethyl-*p*-toluidine as an accelerator. Ultrathin sections (200 nm) were cut and collected on 200-mesh copper finder grids. Fluorescence images of 120-nm sections, from 3 different animals, were acquired using an epifluorescence microscope (Axioskop 2 Plus, Zeiss) equipped with an EMCCD camera (CoolSNAP, Photometrics) before electron microscopy observation.

#### FIB–SEM sample preparation and imaging

Worms were transferred into M9 buffer supplemented with 20% bovine serum albumin (BSA; Sigma-Aldrich, A7030) and loaded into 200 µm-deep flat carriers pre-coated with 1% phosphatidylcholine (Sigma-Aldrich; Leica Biosystems). Samples were cryo-immobilized using an EMPACT-2 high-pressure freezer (Leica Microsystems, Vienna, Austria), as previously described (3). Freeze substitution was performed using an Automated Freeze Substitution system (AFS2; Leica Microsystems, Wetzlar, Germany). Flat carriers were transferred into the AFS2 chamber, pre-cooled to −90 °C, and immersed in a freeze-substitution medium containing acetone with 2% osmium tetroxide and 0.5% uranyl acetate. Samples were maintained at −90 °C for 60–80 h, then warmed to −60 °C at 2 °C h⁻¹ and incubated for 12 h. The temperature was subsequently increased to −30 °C at the same rate, and the samples were incubated for an additional 12 h. Samples were then rapidly warmed to 0 °C, held for 1 h, and then rapidly returned to −30 °C, and incubated for a further 1 h. During this period, the freeze-substitution medium was replaced with anhydrous acetone through three successive 15-min washes. Temperature was then increased from −30 °C to −10 °C at 10 °C h⁻¹. During this step, samples were removed from the AFS and acetone was sequentially replaced by 4% osmium tetroxide (15 min), anhydrous acetone (15 min), 2% uranyl acetate (15 min), and anhydrous acetone (15 min). Samples were then returned to the AFS for resin infiltration. Resin infiltration was performed stepwise using anhydrous Araldite resin (Araldite 502 Epoxy Resin Kit; EMS #13900). Samples were incubated in 30% Araldite in acetone for 1 h at −10 °C, followed by incubation in 50% Araldite during warming from −10 °C to 10 °C (10 °C h⁻¹), and then in 70% Araldite during warming from 10 °C to 20 °C (5 °C h⁻¹). Samples were subsequently incubated for at least 16 h at 20 °C in 100% anhydrous Araldite resin before removal from the AFS. Final embedding was performed in 100% anhydrous Araldite resin supplemented with BDMA accelerator, followed by polymerization at 65 °C for 48 h. Samples were imaged using a Dual Crossbeam 550 microscope (Zeiss). Imaging was performed on a defined volume of approximately 50 µm in width, 30 µm in height, and 20 µm in depth. The acquisition lasted 18 h, yielding a dataset of 1,479 serial images. Images were acquired at a spatial isotropic resolution of 10 nm. Scanning electron microscopy was performed using an accelerating voltage of 1.4 kV and a probe current of 500 pA, with signal detection using SE2 and InLens detectors. Focused ion beam milling was conducted using a current of 1.5 nA. The resulting dataset had a total file size of 2.72 GB. Following the acquisition, the image stack was cropped and initially processed using ATLAS 5, after which alignment between z-slices was corrected using AMST (5). Then, imaging artifacts, including curtaining, non-homogeneous illumination, and uneven z-slicing, were corrected using PyStack3D (6). Image segmentation was performed using Dragonfly (Object Research Systems). Before segmentation, image volumes were cropped to the region of interest using the *ROI Painter* module with the *round brush* tool. Segmentation of the different cellular structures was carried out manually, including the outer membrane, multi-membrane structures, lipid droplets, and the endoplasmic reticulum. Three-dimensional renderings and movies were generated in Dragonfly using the *Movie Maker* module. This study included FIB-SEM data from three *pdzd-8* mutants.

#### RNA interference

Standard NGM plates were modified with the addition of 2 mM final isopropyl-β-d-thiogalactopyranoside (IPTG) and 25 μg/ml final carbenicillin. The RNAi experiments were performed using the feeding procedure (7). Feeding RNAi clones were purchased from Open Biosystem. 30-40 L4 stage *C. elegans* were picked to each plate seeded with HT115 (DE3) bacteria carrying double-strand RNA expression constructs for either ctrl L4440 (the empty vector L4440 (pPD129.36)), *lgg-1* (C32D5.9), *atg-3*, *epg-5* (M7.5) or *rab-7* (ZK593.6) at 20°C. HT115 (DE3) bacteria carrying the empty vector alone (L4440) were used as a negative control for all experiments. When worms reached adulthood, they were transferred to fresh RNAi plates and allowed to lay eggs for 2 hours to generate synchronized F1 self-progeny, which was further cultured to the early L4 stage.

#### Phenotype analysis

Microscope analysis, imaging, and phenotype analysis were conducted on synchronized worms. Synchronization of the worms was achieved by picking 30-50 young gravid adults from the original plate and transferring them to a fresh plate, where they were allowed to lay eggs for one hour at 20°C. Subsequently, the adults were removed, and the plates were maintained at 20°C to permit embryo hatching and larval development until the designated time points for imaging, developmental phenotype analysis, or acute heat stress. The early L4 larval stage was confirmed under DIC (differential interference contrast) imaging in a compound microscope, as evidenced by the structure of the elongated gonad arms.

To measure body size, mutant and control worms were selected and mounted on slides in M9 solution containing 40 mM NaN3. Thereafter, photographs were taken under Nomarski optics using a 10X objective (Zeiss Axiovert). Subsequently, the worm body length was determined using ImageJ.

Life-span assays were conducted with synchronised L4 larvae. The worms were then transferred to NGM agar plates containing OP50 bacteria. Subsequently, 150 young adult animals, collected from the aforementioned plates, were distributed evenly across 15 fresh NGM plates seeded with bacteria. To prevent contamination between the test worms and their progeny, the worms were transferred to a new plate daily until they ceased egg production, after which they were transferred every 2 days. The number of dead and live worms was recorded daily. The animals were designated as deceased when they failed to respond to repeated gentle tactile stimulation with a platinum wire. The missing worms were scored as censored data. The lifespan is defined in days, commencing with the transfer of young adults to plates (adult lifespan = 0) and concluding with the scoring of mortality. The lifespan is expressed as the mean ± S.E.M. (standard error of the mean). The experiments were conducted in triplicate at 20 °C. For aHS, synchronized early L4-stage larvae were incubated at 37 °C for 60 minutes in an incubator (Binder).

#### Image analyses

All images were processed using ImageJ/Fiji (8). In some experiments, the fluorescent signal identification and quantification were performed using a macro that included the Trainable Weka Segmentation (TWS) plugin. For segmentation, too or three classes were defined: fluorescent signals of interest and background. Manual labeling was performed by selecting representative regions of interest (ROIs) for each class using the polygonal selection tool. Following labeling, the classifier was trained on the selected features. Upon completion of training, the classifier was applied to the entire image to segment fluorescent signals from the background. The segmented image (32 bits) was then converted to binary using the automatic threshold method: intermodes.

Fluorescent signals of interest were quantified using the ’Analyze Particles’ function in ImageJ, with size and circularity exclusion thresholds applied to differentiate between noise and valid signals. The output included the total particle count and signal area. The ROIs from the ’Analyze Particles’ function were overlaid on the raw images to inspect the consistency of the results visually. The classifier was saved and applied by the macro to additional images to ensure uniformity in batch processing. Results were confirmed by visual inspection of the segmented images, and the data were subsequently exported for statistical analysis.

#### Colocalization experiments

Quantification of colocalization between GFP and mCherry signals was conducted using ImageJ (NIH) with the JACOP plugin (9). Before analysis, background fluorescence was subtracted from each channel with a threshold. Manders’ colocalization coefficients (M1 and M2) were calculated to quantify the degree of colocalization between the two fluorophores. M1 represents the fraction of GFP signal overlapping with mCherry, and M2 represents the fraction of mCherry signal overlapping with GFP. The coefficients range from 0 (no colocalization) to 1 (complete colocalization). For statistical analysis, Manders’ coefficients from three independent experiments were calculated and expressed as mean ± standard deviation (SD).

#### Statistical Analysis

All experiments were performed in at least three independent biological replicates. Data were expressed as mean ± standard deviation (SD) unless otherwise noted. Statistical analysis was performed using GraphPad Prism 9 or the R package (https://www.R-project.org/). The boxplots display the first quartile (Q1/25th percentile), the median (Q2/50th percentile), and the third quartile (Q3/75th percentile). The whiskers extend up to 1.5 times the interquartile range. Before statistical testing, data were assessed for normality using a QQ plot (quantile-quantile plot) and for homogeneity of variances using Levene’s test. Data that followed a normal distribution were analyzed using parametric tests, while non-normally distributed data were analyzed using non-parametric tests. For comparisons between two groups, a two-tailed Student’s t-test was used for normally distributed data with homogeneity of variances; otherwise, a Welsh test was used. When the assumption of normality was not valid, the Mann-Whitney U test was used. Statistical significance is indicated in the figures with asterisks as follows: p < 0.05 (*), p < 0.01 (**), p < 0.001 (***), and p < 0.0001 (****). All statistical tests and figures were generated using GraphPad Prism or the R package.

**SI Fig. S1 - related to Figure 1.**
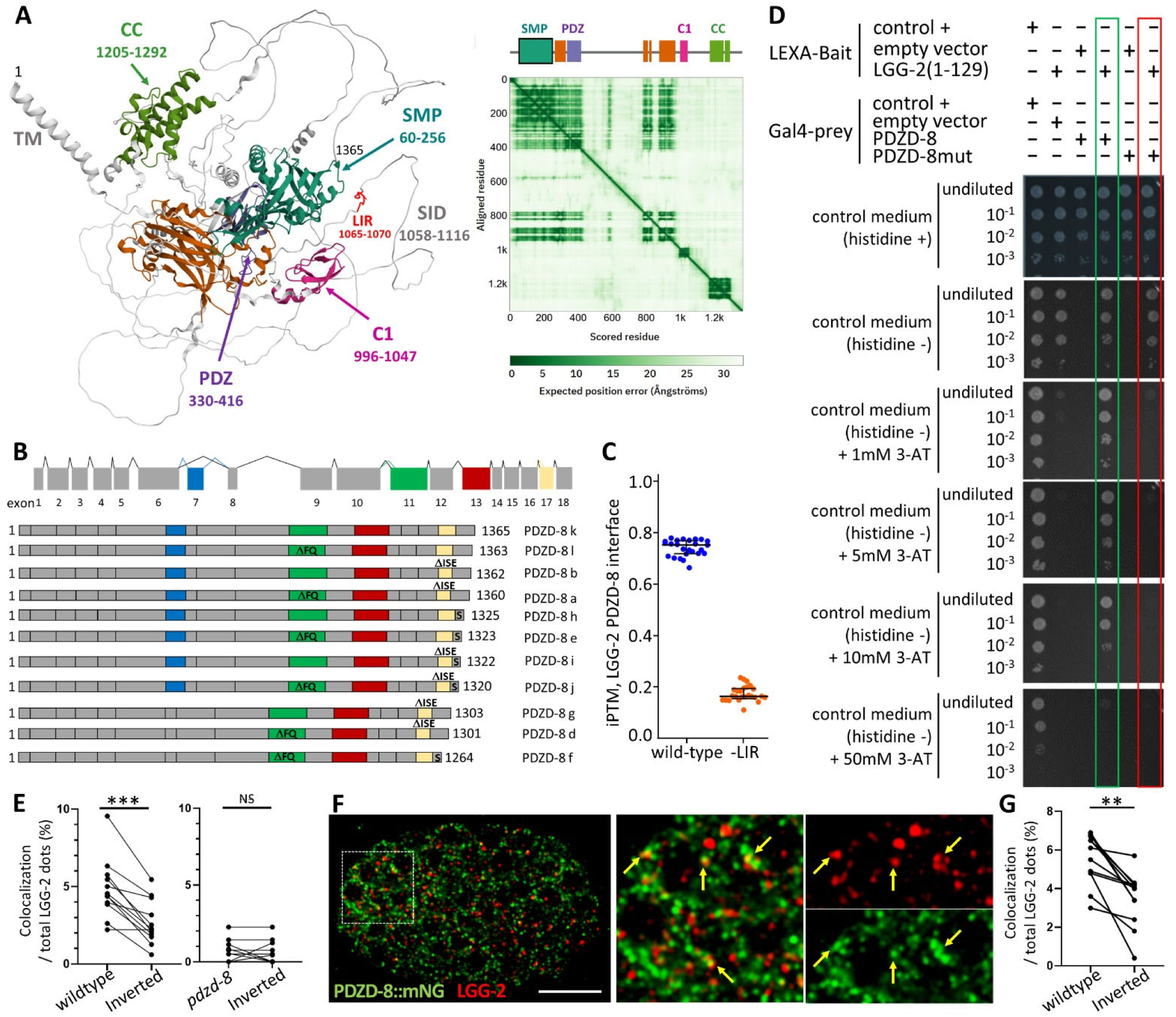
(A) PDZD-8 3D-structure from AlphaFold Protein Structure Database (AF-A0A486WTK6-F1-v6) and Predicted Aligned Error plot showing the modelization of TM (Trans Membrane domain), SMP (Synaptotagmin-like mitochondrial-lipid-binding), PDZ, C1, CC (coiled coil) domains. (B) Schematic of the *C. elegans pdzd-8* genetic locus. Alternative transcription initiation, termination, and splicing result in 13 transcripts encoding 11 protein isoforms. (C) Boxplots of interface ipTM for the LGG2 – PDZD-8 with or without the LIR motif. n= 25. Boxes show the median and interquartile range (IQR); whiskers show 1.5×IQR. (D) Yeast two-hybrid assays confirmed a direct interaction between PDZD-8 and LGG-2. The inhibitor 3-aminotriazole (3AT) is used to assess the strength of the interactions and block the auto-activation of LGG-2. One of two independent experiments using different clones is shown. (E) LGG-2 and PDZD-8 partially colocalize in the embryo. Quantification of GFP::LGG-2 and PDZD-8 colocalization, based on distance between centers of mass, in both wild-type and *pdzd-8(syb3082)*. Inverted indicates the random colocalization after the original red channel was rotated to 180°. (F-G) Single merged confocal images of a PDZD-8::mNG and LGG-2 embryos immunostained with anti-mNG (green) and anti-LGG-2 (red) (F) and quantification (similar to E) G. Scale bar is 10µm.Kruskal-Wallis test with Bonferroni correction ** p<0.01 *** p<0.001 NS: non-significant.

**SI Fig. S2 - related to Figure 1 and Figure 2.**
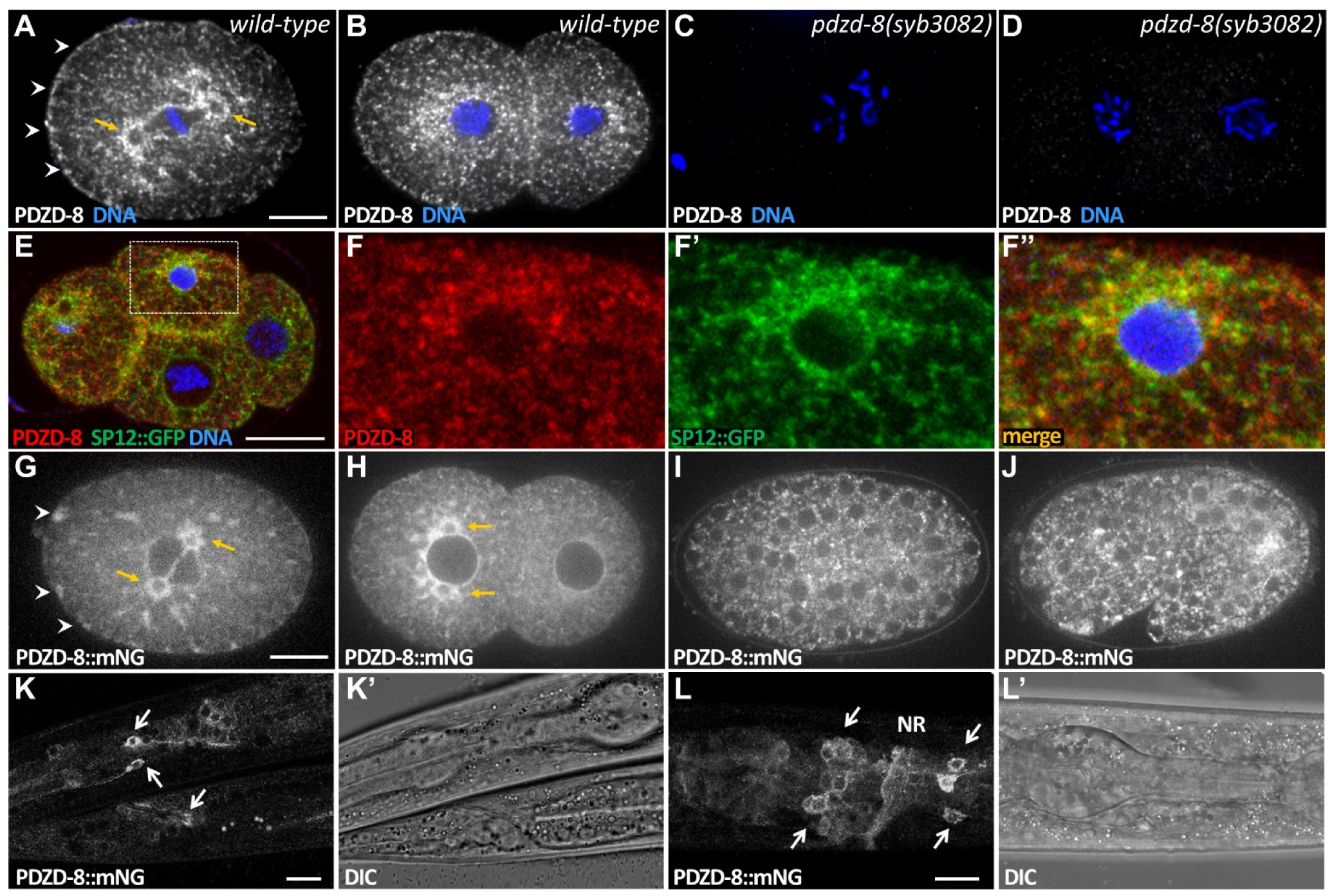
PDZD-8 expression pattern and subcellular localization (A-D) Anti-PDZD-8 antibody validation experiment. Confocal images of the equatorial plane of 2-cells and 4-cells embryos from control (A-B) and pdzd-8(syb3082) null mutant (C-D), immunolabelled with PDZD-8 antibody (white) and stained with Hoechst (blue). In early embryos, PDZD-8 is predominantly detected in the perinuclear and cortical endoplasmic reticulum (ER), as well as the centriculum (ER surrounding the centrosome). (E-F) Confocal images of a wild-type 4-cell embryo (mid-plane), showing the colocalization of PDZD-8 with the ER protein SP-12::GFP. The embryo is immunolabeled with anti-PDZD-8 (red) and anti-GFP (green) antibodies. Panels F–F″ are magnified views of the box in E. (G-J) Spinning disc confocal images showing the mid-plane 1-cell (G), 2-cells (H), 100-cells (I), and 1.5-fold (J) embryo expressing PDZD::mNeonGreen (mNG). In early embryos, the PDZD-8::mNG pattern is similar to endogenous PDZD-8. It is weakly expressed early in embryogenesis and throughout the C. elegans life cycle, and is enriched in nerves and the epidermis. The scale bar is 10 µm.

**SI Fig. S3 - related to Figure2.**
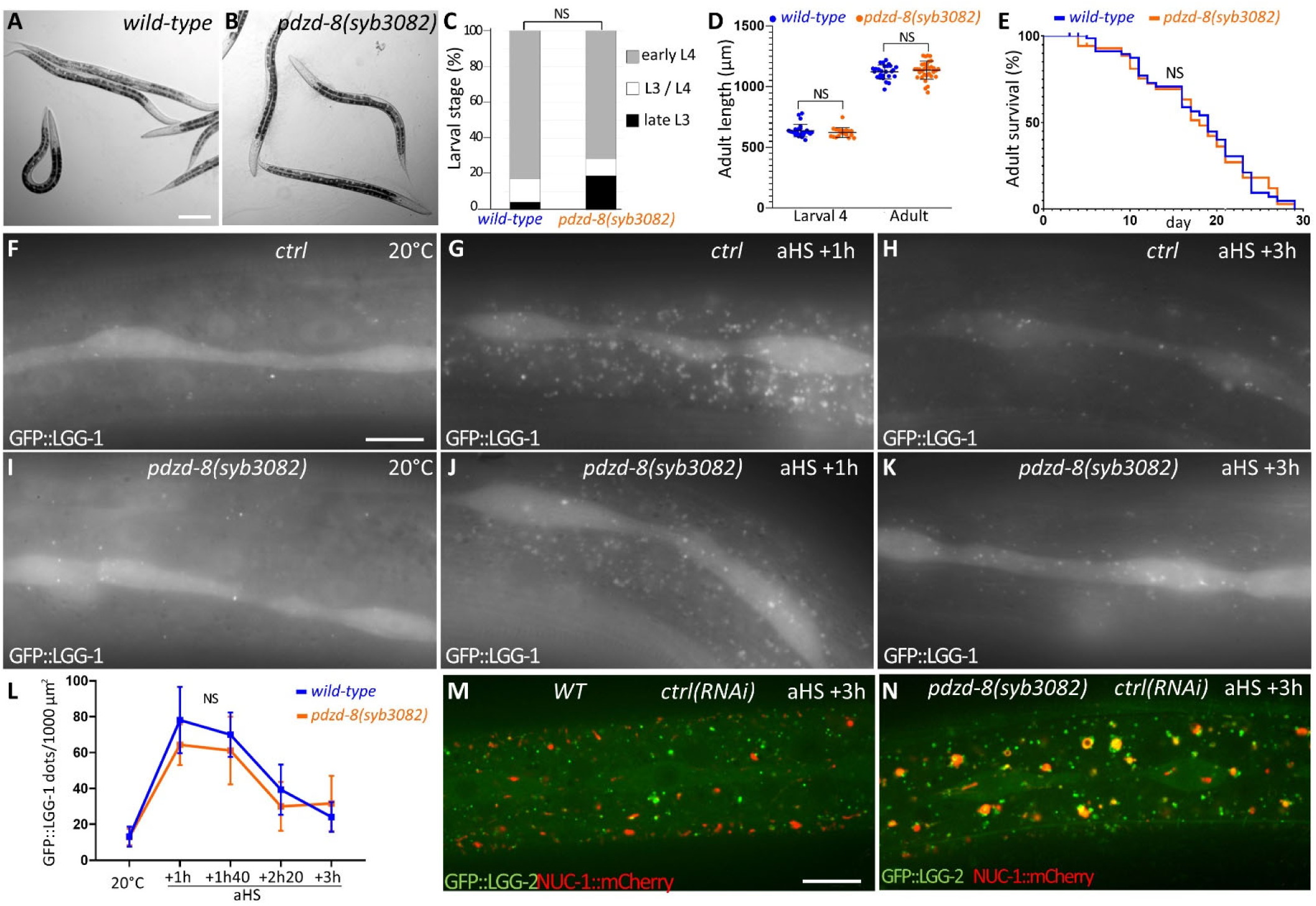
*pdzd-8(syb3082)* mutant has no developmental delay nor shorter lifespan. (A-E) Early L4 worms from both control (A) and *pdzd-8(syb3082),* illustrating rather homogenous and similar morphology and size. (C) Histograms of late L3 (lL3), transition L3/L4 and early L4 in wild-type and *pdzd-8(syb3082*) larvae, 43 hours post-hatch at 20°C. (D) Boxplots display the quantification of body size for wild-type and *pdzd-8* mutant animals at the L4 larval and adult stages. Bars are mean and SD. n animals> 19. Mann-Whitney U test, NS: no significant difference. (E) Lifespan analysis of wild-type (N2), *pdzd-8* (syb3082) animals (one representative experiment from 3 independent replicates). n animals > 100, log-rank test, NS: no significant. (F-L) Wide-field images of GFP::LGG-1 in the epidermis of control and *pdzd-8(syb3082)* mutant L4 animals at 20°C (F, I) or 1h (G, J) and 3h after aHS (H, K). (L) Quantification of GFP::LGG-1 structures showing no significant difference (NS). n animals = x, Welch t-test. Representative data from 3 replicates. (M, N) controls for Figure 2 (R-W). Confocal images of GFP::LGG-2 and NUC-1::mCherry 3h post aHS in the epidermis of control and *pdzd-8 (syb3082)* animals submitted to an empty RNAi vector (L4440) as control. The scale bars are 100 µm (A, B) and 10 µm (F-N).

**SI Fig. S4 - related to Figure3.**
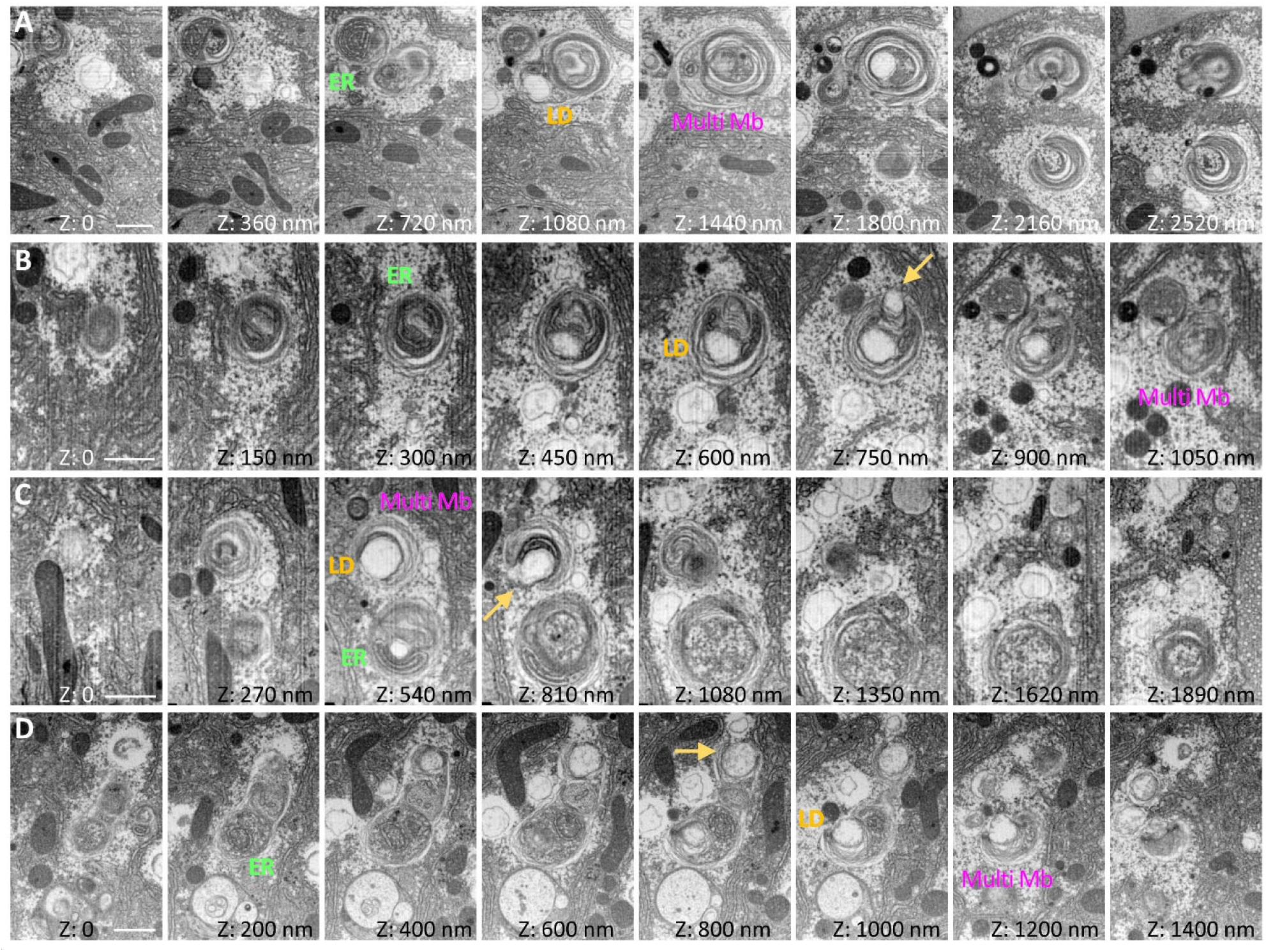
Lipophagy is induced by acute heat stress. (A-D) Focused ion beam–scanning electron microscopy (FIB-SEM) imaging of autolysosome clusters in the epidermis of *pdzd-8(syb3082)* mutants 3h after aHS. Eight panels have been extracted from the image series, and the Z position is indicated for each. A-C are zooms from the same FIB-SEM series (resolution 10 nm) and D from another FIB-SEM on a different worm (resolution 10 nm). FIB-SEM reveals the presence of packed ER, membranous whorls (Multi Mb), and undegraded lipid droplets (LD) inside the autolysosome, suggesting that the aHS induces lipophagy in the epidermis. Yellow arrows indicate partially engulfed LD, which may indicate microlipophagy events. The scale bar is 1µm.

**SI Fig. S5 - related to Figure 4.**
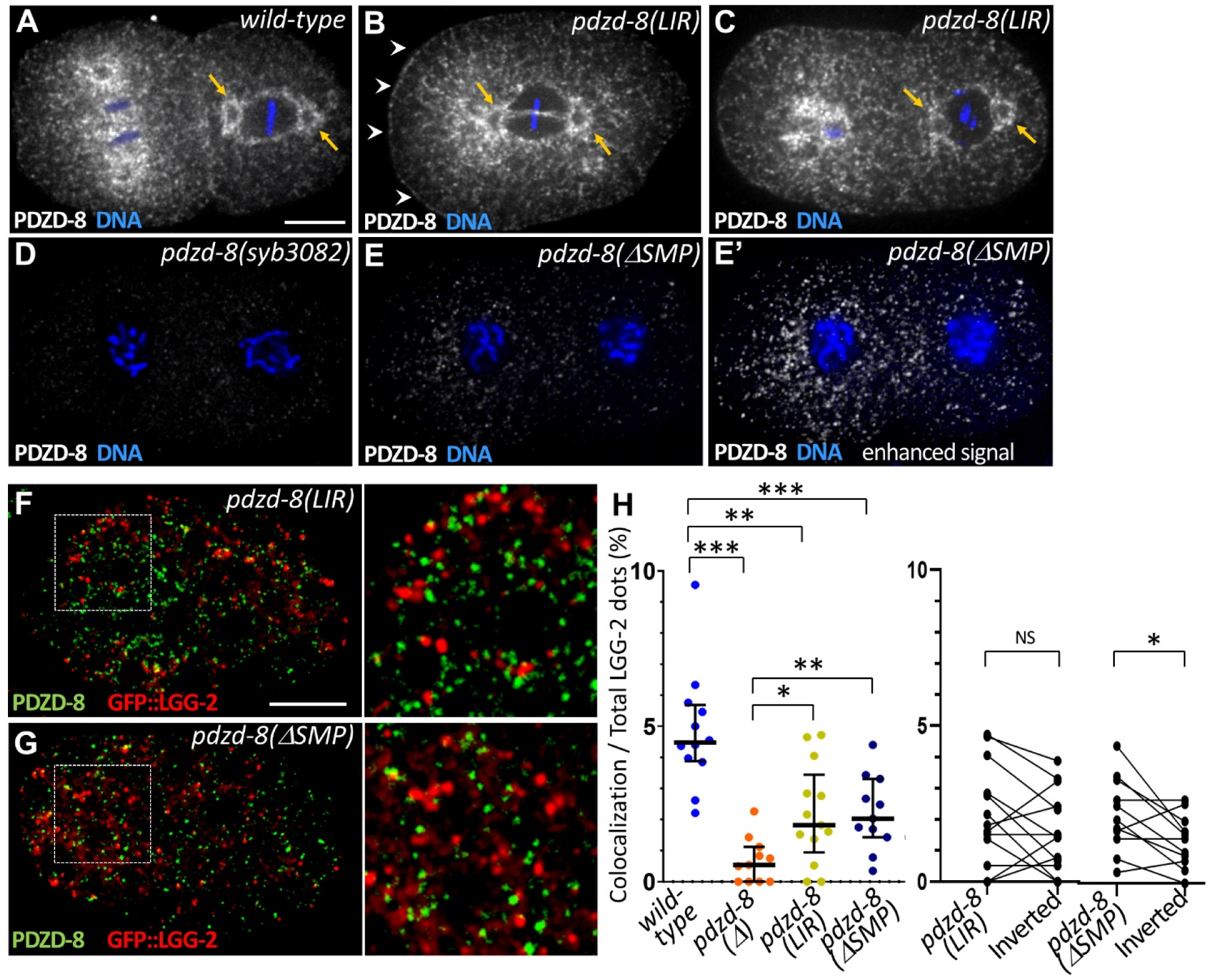
The LIR motif is essential for PDZD-8 colocalization with LGG-2. (A-E) Confocal images of the equatorial plane of 2-cell embryos from control (A), *pdzd-8(LIR)*(B, C), *pdzd-8(syb3082)*(D), and *pdzd-8(ΔSMP)*(E) mutants immunolabelled with PDZD-8 antibody (white) and stained with Hoechst (blue). White arrowheads point to cortical ER, and orange arrows indicate the centriculum, an ER compartment surrounding the centrosome. PDZD-8 subcellular distribution is preserved in *pdzd-8(LIR)* mutant, but markedly reduced in *pdzd-8(ΔSMP)*. (F-H) Colocalization of PDZD-8 with LGG 2 is altered in LIR and SMP mutants. Single merged confocal images of a GFP::LGG-2 embryo, immunostained with antiPDZD-8 (green) and antiGFP (red) antibodies in *pdzd-8(LIR)*(F), and *pdzd-8(ΔSMP)*(G) mutants. The right panels show magnified views of the box regions. Quantification of colocalization based on the distance between centers of mass. Inverted indicates the random colocalization after the original red channel was rotated to 180°. Bars indicate the median and the 25th and 75th percentiles. Kruskal-Wallis test with Bonferroni correction ***p<0.001, **p<0.01, *p<0.05, NS: non-significant. The scale bar is 10 µm.

**Movie S1 (separate file).**

Time-lapse spinning-disk confocal live imaging of one epidermal field of a *pdzd-8(syb3082)* mutant animal acquired every 15 minutes over a 2h45 period, 3h post aHS. Lysosomes (NUC-1::mCherry-labeled) appear as large red puncta, and autophagosomes (GFP::LGG-2 positive) as small green puncta. The scale bar is 1 µm.

**Movie S2 (separate file).**

FIB-SEM imaging series corresponding to Figure 3 (D) of autolysosome clusters in the epidermis of *pdzd-8(syb3082)* 3h post aHS. (resolution 10nm). The scale bar is 1 µm.

**Movie S3 (separate file).**

A 3D model of an enlarged autolysosome in the epidermis of *pdzd-8(syb3082)* mutant 3h post aHS. The model was obtained from the FIB-SEM imaging series shown in Movie S2, corresponding to Figure 3 (D). The external membrane of the autolysosome is in red. Internal compartments are the membranous whorls (blue), packed ER (green), and lipid droplets (yellow).

## References

1. M. Ebner, et al., Nutrient-regulated control of lysosome function by signaling lipid conversion. Cell 186, 5328–5346.e26 (2023).

2. H. Liu, et al., PtdIns4P exchange at endoplasmic reticulum-autolysosome contacts is essential for autophagy and neuronal homeostasis. Autophagy 19, 2682–2701 (2023).

3. M. Ebner, F. Fröhlich, V. Haucke, Mechanisms and functions of lysosomal lipid homeostasis. Cell Chem Biol 32, 392–407 (2025).

4. C. Yang, X. Wang, Lysosome biogenesis: Regulation and functions. J Cell Biol 220, e202102001 (2021).

5. W. W.-Y. Yim, N. Mizushima, Lysosome biology in autophagy. Cell Discov 6, 6 (2020).

6. H.-M. Shen, N. Mizushima, At the end of the autophagic road: an emerging understanding of lysosomal functions in autophagy. Trends Biochem Sci 39, 61–71 (2014).

7. T. N. Nguyen, et al., Atg8 family LC3/GABARAP proteins are crucial for autophagosome-lysosome fusion but not autophagosome formation during PINK1/Parkin mitophagy and starvation. J Cell Biol 215, 857–874 (2016).

8. H. Wang, et al., GABARAPs regulate PI4P-dependent autophagosome:lysosome fusion. Proc Natl Acad Sci U S A 112, 7015–7020 (2015).

9. M. Manil-Ségalen, et al., The C. elegans LC3 acts downstream of GABARAP to degrade autophagosomes by interacting with the HOPS subunit VPS39. Dev. Cell 28, 43–55 (2014).

10. T. Baba, D. J. Toth, N. Sengupta, Y. J. Kim, T. Balla, Phosphatidylinositol 4,5-bisphosphate controls Rab7 and PLEKHM1 membrane cycling during autophagosome-lysosome fusion. EMBO J 38, e100312 (2019).

11. H. Jeong, J. Park, Y. Jun, C. Lee, Crystal structures of Mmm1 and Mdm12-Mmm1 reveal mechanistic insight into phospholipid trafficking at ER-mitochondria contact sites. Proc Natl Acad Sci U S A 114, E9502–E9511 (2017).

12. Y. Elbaz-Alon, et al., PDZD8 interacts with Protrudin and Rab7 at ER-late endosome membrane contact sites associated with mitochondria. Nat Commun 11, 3645 (2020).

13. Y. Gao, J. Xiong, Q.-Z. Chu, W.-K. Ji, PDZD8-mediated lipid transfer at contacts between the ER and late endosomes/lysosomes is required for neurite outgrowth. J Cell Sci 135, jcs255026 (2022).

14. H. Khan, L. Chen, L. Tan, Y. J. Im, Structural basis of human PDZD8-Rab7 interaction for the ER-late endosome tethering. Sci Rep 11, 18859 (2021).

15. A. Guillén-Samander, X. Bian, P. De Camilli, PDZD8 mediates a Rab7-dependent interaction of the ER with late endosomes and lysosomes. Proc. Natl. Acad. Sci. U.S.A. 116, 22619–22623 (2019).

16. Y. Hirabayashi, et al., ER-mitochondria tethering by PDZD8 regulates Ca2+ dynamics in mammalian neurons. Science 358, 623–630 (2017).

17. K. Nakamura, et al., Mitochondrial complexity is regulated at ER-mitochondria contact sites via PDZD8-FKBP8 tethering. Nat Commun 16, 3401 (2025).

18. M. Shirane, et al., Protrudin and PDZD8 contribute to neuronal integrity by promoting lipid extraction required for endosome maturation. Nat Commun 11, 4576 (2020).

19. D. Jeyasimman, et al., PDZD-8 and TEX-2 regulate endosomal PI(4,5)P2 homeostasis via lipid transport to promote embryogenesis in C. elegans. Nat Commun 12, 6065 (2021).

20. H. Yang, et al., LYVAC/PDZD8 is a lysosomal vacuolator. Science 389, eadz0972 (2025).

21. K. Morita, et al., PDZD8-deficient mice accumulate cholesteryl esters in the brain as a result of impaired lipophagy. iScience 25, 105612 (2022).

22. R. S. Thakur, K. M. O’Connor-Giles, PDZD8 promotes autophagy at ER-lysosome membrane contact sites to regulate activity-dependent synaptic growth. Cell Rep 44, 115483 (2025).

23. C. Serot, et al., Reticulon-dependent ER-phagy mediates adaptation to heat stress in C. elegans. Curr Biol 35, 2365–2378.e7 (2025).

24. C. Hoffmann, et al., Membrane-protein-mediated phase separation orchestrates organelle contact sites. Mol Cell 86, 135–149.e9 (2026).

25. J. Jumper, et al., Highly accurate protein structure prediction with AlphaFold. Nature 596, 583–589 (2021).

26. F. Wu, et al., Structural Basis of the Differential Function of the Two C. elegans Atg8 Homologs, LGG-1 and LGG-2, in Autophagy. Mol Cell 60, 914–929 (2015).

27. R. Maheshwari, et al., A membrane reticulum, the centriculum, affects centrosome size and function in Caenorhabditis elegans. Curr Biol 33, 791–806.e7 (2023).

28. Y. Chen, et al., Autophagy facilitates mitochondrial rebuilding after acute heat stress via a DRP-1-dependent process. J Cell Biol 220 (2021).

29. A. Meléndez, et al., Autophagy genes are essential for dauer development and life-span extension in C. elegans. Science 301, 1387–1391 (2003).

30. A. Alberti, X. Michelet, A. Djeddi, R. Legouis, The autophagosomal protein LGG-2 acts synergistically with LGG-1 in dauer formation and longevity in C. elegans. Autophagy 6, 622–633 (2010).

31. I. G. Ganley, P.-M. Wong, N. Gammoh, X. Jiang, Distinct autophagosomal-lysosomal fusion mechanism revealed by thapsigargin-induced autophagy arrest. Mol Cell 42, 731–743 (2011).

32. K. Hegedűs, et al., The Ccz1-Mon1-Rab7 module and Rab5 control distinct steps of autophagy. Mol Biol Cell 27, 3132–3142 (2016).

33. A. H. Al-Amri, et al., PDZD8 Disruption Causes Cognitive Impairment in Humans, Mice, and Fruit Flies. Biol Psychiatry 92, 323–334 (2022).

34. R. A. Bharadwaj, et al., Genetic risk mechanisms of posttraumatic stress disorder in the human brain. J Neurosci Res 96, 21–30 (2018).

35. A. D. Pantiru, et al., Autistic behavior is a common outcome of biallelic disruption of PDZD8 in humans and mice. Mol Autism 16, 14 (2025).

36. D. S. Grunwald, N. M. Otto, J.-M. Park, D. Song, D.-H. Kim, GABARAPs and LC3s have opposite roles in regulating ULK1 for autophagy induction. Autophagy 16, 600–614 (2020).

37. J. Zhao, Z. Li, J. Li, The crystal structure of the FAM134B-GABARAP complex provides mechanistic insights into the selective binding of FAM134 to the GABARAP subfamily. FEBS Open Bio 12, 320–331 (2022).

38. S. Nakamura, T. Yoshimori, New insights into autophagosome-lysosome fusion. J Cell Sci 130, 1209–1216 (2017).

39. C. Zhou, et al., Recycling of autophagosomal components from autolysosomes by the recycler complex. Nat Cell Biol 24, 497–512 (2022).

40. S. C. Li, et al., The signaling lipid PI(3,5)P₂ stabilizes V₁-V(o) sector interactions and activates the V-ATPase. Mol Biol Cell 25, 1251–1262 (2014).

## SI References

1. S. Brenner, The genetics of Caenorhabditis elegans. Genetics 77, 71–94 (1974).

2. J. Jumper, et al., Highly accurate protein structure prediction with AlphaFold. Nature 596, 583–589 (2021).

3. C. Largeau, R. Legouis, Correlative Light and Electron Microscopy to Analyze LC3 Proteins in Caenorhabditis elegans Embryo. Methods Mol Biol 1880, 281–293 (2019).

4. C. Largeau, E. Culetto, R. Legouis, Subcellular Localization of ESCRT-II in the Nematode C. elegans by Correlative Light Electron Microscopy. Methods Mol Biol 1998, 49–61 (2019).

5. J. Hennies, et al., AMST: Alignment to Median Smoothed Template for Focused Ion Beam Scanning Electron Microscopy Image Stacks. Sci Rep 10, 2004 (2020).

6. Quéméré & David. (2024). PyStack3D: A python package for fast image stack correction. Journal of Open Source Software, 9(101), 7079

7. R. S. Kamath, J. Ahringer, Genome-wide RNAi screening in Caenorhabditis elegans. Methods 30, 313–321 (2003).

8. J. Schindelin, et al., Fiji: an open-source platform for biological-image analysis. Nat Methods 9, 676–682 (2012).

9. S. Bolte, F. P. Cordelières, A guided tour into subcellular colocalization analysis in light microscopy. J Microsc 224, 213–232 (2006).

